# Kv3.3 subunits control presynaptic action potential waveform and neurotransmitter release at a central excitatory synapse

**DOI:** 10.1101/2021.11.02.466934

**Authors:** Amy Richardson, Victoria Ciampani, Mihai Stancu, Sherylanne Newton, Joern R. Steinert, Nadia Pilati, Bruce P. Graham, Conny Kopp-Scheinpflug, Ian D. Forsythe

## Abstract

Kv3 potassium currents mediate rapid repolarization of action potentials (AP), supporting fast spikes and high repetition rates. Of the four Kv3 gene family members, Kv3.1 and Kv3.3 are highly expressed in the auditory brainstem and we exploited this to test for subunit-specific roles at the calyx of Held presynaptic terminal. Deletion of Kv3.3 (but not Kv3.1) increased presynaptic AP duration and facilitated transmitter release, which in turn enhanced short-term depression during high frequency transmission. The response to sound was delayed in the Kv3.3KO, with higher spontaneous and lower evoked firing, thereby reducing signal-to-noise ratio. Computational modelling showed that the enhanced EPSC and short-term depression in the Kv3.3KO reflected increased vesicle release probability and accelerated activity-dependent vesicle replenishment. We conclude that Kv3.3 is the presynaptic ‘delayed rectifier’, enabling short duration, precisely timed APs to maintain transmission at high frequencies and during sustained synaptic activity.

## Introduction

Kv3 voltage-gated potassium currents rapidly repolarize action potentials and underlie fast-spiking neuronal phenotypes, enabling high frequency firing with temporal precision (Rudy & McBain, 2001; Kaczmarek & Zhang, 2017). Kv3 channels are expressed throughout the brain including the hippocampus, cortex, cerebellum and auditory brainstem (Weiser ***et al.***, 1994; Du ***et al.***, 2000; Lien & Jonas, 2003). These precisely located Kv3 channels influence dendritic integration (Zagha ***et al.***, 2010) and somatic action potential waveform (Espinosa ***et al.***, 2008; Rowan ***et al.***, 2014; Choudhury ***et al.***, 2020). They are also located in some axons, including nodes of Ranvier (Devaux ***et al.***, 2003) and synaptic terminals where they modulate neurotransmitter release (Forsythe, 1994; Wang ***et al.***, 1998; Ritzau-Jost ***et al.***, 2021).

There are four Kv3 genes (kcnc1-4) specifying alpha subunits (Kv3.1-3.4) that assemble as tetramers (Coetzee ***et al.***, 1999; Rudy ***et al.***, 2002).The transmembrane domains are generally well conserved across subunits but both the N- and C-terminal domains differ, creating distinct activation and inactivation properties, in addition to independent interactions with cytoskeletal proteins (Zhang ***et al.***, 2016; Stevens ***et al.***, 2021) and regulation by phosphorylation (Song ***et al.***, 2005; Steinert ***et al.***, 2011). Recombinant Kv3 channels display ultra-fast kinetics (Grissmer ***et al.***, 1994; Labro ***et al.***, 2015) but it is often difficult to reconcile their properties with ‘native’ Kv3 channels where multiple subunit genes are often expressed in a specific neuron (Johnston ***et al.***, 2010). Transgenic knockouts of one subunit generally show mild phenotypes, consistent with heterogeneous composition of native channels and functional redundancy (Joho ***et al.***, 1999; Espinosa ***et al.***, 2001; Joho ***et al.***, 2006). Co-expression of Kv3.1 and Kv3.3 has been widely observed in different brain regions (Chang ***et al.***, 2007a) and we showed that Kv3.1 and Kv3.3 could compensate for each other in principal neurons of the medial nucleus of the trapezoid body (MNTB) within the auditory brainstem, maintaining high frequency somatic action potential firing (Choudhury ***et al.***, 2020).

Changes in AP waveform at the synaptic terminal critically control calcium influx and neurotransmitter release (Borst & Sakmann, 1998; Forsythe ***et al.***, 1998; Yang ***et al.***, 2014), this in turn influences short-term plasticity (Sakaba & Neher, 2003; Wong ***et al.***, 2003; Hennig ***et al.***, 2008; Neher, 2017) and presynaptic forms of long term potentiation at mossy fibre terminals (Geiger & Jonas, 2000). In the present study we took advantage of the auditory brainstem which has a high density of Kv3 channels to investigate the role of Kv3 subunits at the presynaptic terminal. The binaural auditory nuclei must rapidly integrate AP trains transmitted from left and right cochlea with microsecond accuracy (Beiderbeck ***et al.***, 2018; Joris & Trussell, 2018). The region expresses Kv3.1 and Kv3.3 subunits, with little or no Kv3.2 and Kv3.4 (Choudhury ***et al.***, 2020). This system is operating at the ‘biophysical limit’ of information processing, in that fast conducting axons and giant synapses with nano-domain localization of P/Q Ca^2+^ channels, combine with postsynaptic expression of fast AMPARs, short membrane time-constants and exceptionally rapid APs to enable binaural sound localization (Schneggenburger & Forsythe, 2006; Johnston ***et al.***, 2010; Young & Veeraraghavan, 2021). In this study we test whether Kv3.1 or Kv3.3 subunits have a specific presynaptic role by examining the calyx of Held AP waveform and transmitter release on deletion of one Kv3 subunit. We then demonstrate the *in vivo* consequences for brainstem auditory processing of sound and conclude that the Kv3.3 subunit dominates fast AP repolarization at this excitatory synapse.

## Results

The experiments reported here were conducted using *in vitro* brainstem slices from CBA/CaL mice or transgenic mice backcrossed onto CBA/CaL that lacked either Kv3.3 or Kv3.1 (the genotypes are referred to as WT, Kv3.3KO and Kv3.1KO). The influence of these deletions was also assessed *in vivo* using extracellular recording from the MNTB. Whole cell patch clamp recording from the calyx of Held and MNTB principal neurons was employed to determine the contribution of Kv3 subunits to the presynaptic AP waveform, and to assess the impact on transmitter release. We previously established that Kv3.1 and Kv3.3 mRNA is highly expressed in the MNTB (Choudhury ***et al.***, 2020). Additionally, we conducted mRNA sequencing of the cochlear nucleus and confirmed a similar expression pattern in this brain region containing the globular bushy cells that give rise to the calyx of Held synaptic terminal. The percent contribution of each subunit gene to the cochlear nucleus Kv3 mRNA was 30.3±9.1%, 7.7±2.8%, 52.0±6.0% and 10.0±1.6%, respectively; these values are similar to those measured in the MNTB (Choudhury ***et al.***, 2020) (for data see summary statistics).

### Kv3 channels contribute to action potential repolarization at the calyx of Held terminal

Previous reports have employed potassium channel blockers (4-aminopyridine or tetraethylammonium) to show that Kv3 currents contribute to fast repolarization in postsynaptic APs in the MNTB (Forsythe, 1994; Brew & Forsythe, 1995; Wang ***et al.***, 1998). Low concentrations of extracellular TEA (1 mM) give a relatively selective block of Kv3 potassium channels (Johnston ***et al.***, 2010). Whole terminal voltage-clamp recordings of the calyx terminal revealed an outward current, as shown in the current-voltage relations in Fig. 1A, with 1 mM TEA blocking 28.7% of the total outward current at a command voltage of +10mV (current amplitude in WT= 9.37±2.6 nA, n=7 calyces; WT+TEA= 6.68±0.9 nA, n=6; unpaired t-test, P=0.037; mean ± SD). Injection of depolarizing current steps (100 pA, 50 ms) under current-clamp triggered a single AP in the presynaptic terminal, (Fig. 1B) which increased in duration on perfusion of 1 mM TEA (Fig. 1C; with inset showing ±TEA overlay). In tissue from WT mice the mean AP half-width increased from 0.28±0.02 ms (control; n=9) to 0.51±0.1 ms (n=7) in the presence of 1 mM TEA (Fig. 1E), supporting the hypothesis that presynaptic Kv3 channels are present and contribute to AP repolarization at the calyx of Held.

**Figure 1:**
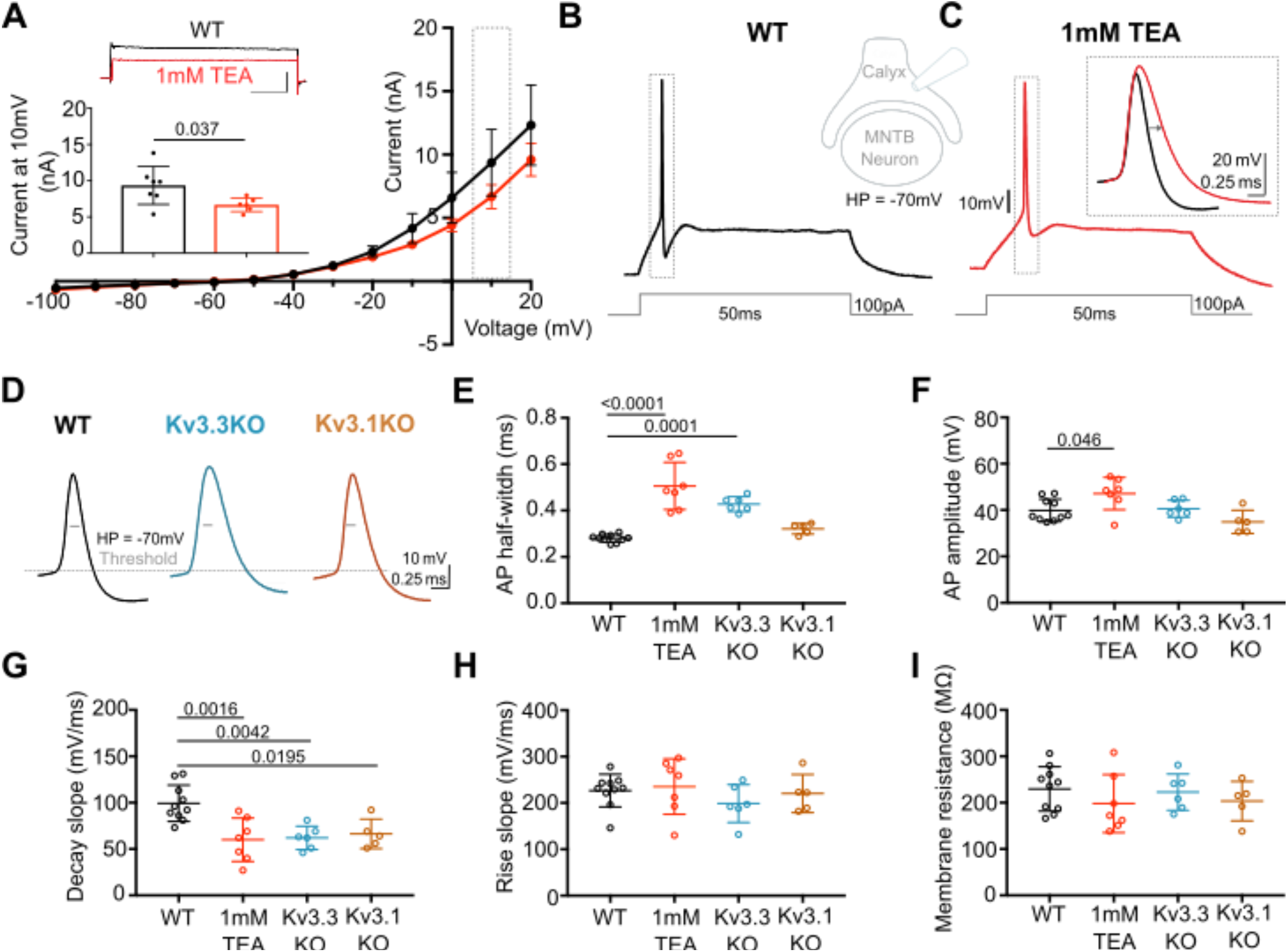
Presynaptic AP duration is increased by TEA or Kv3.3 deletion. **A:** Current-Voltage (I-V) relationship for potassium currents in the calyx of Held terminal of WT mice (P10-P12) in control (black; n=7 terminals, 4 mice) and TEA, 1mM (orange, n=6 terminals, 5 mice; HP= −70mV). **Inset (top):** Example current traces in response to voltage command of +10mV step in WT mouse ± 1 mM TEA. Scale bars = 5 nA and 20ms. **Inset (lower):** Bar graph of mean currents ± SD, measured on step depolarization to +10mV (from HP −70 mV) ±1 mM TEA. Outward Currents are significantly reduced by TEA (student’s t-test, unpaired, P=0.0386). **B:** WT calyx AP (black trace) evoked by 100pA step current injection; **inset** - diagram of recording configuration. **C:** WT calyx AP in the presence of TEA (1 mM, orange trace); inset – overlaid WT APs ± TEA (orange) as indicated by dotted box (grey) around APs in B and C. **D:** Representative AP traces from calyx terminals of WT (black), Kv3.3KO (blue) and Kv3.1KO (orange); double arrows indicate the half-width of WT AP. AP threshold is indicated by the grey dashed line. **E:** AP half-width measured as time difference between rise and decay phases at 50% maximal amplitude. Half-width is significantly increased in TEA and in Kv3.3KO; N is individual terminals: WT N = 9 from 6 animals; TEA = 7 from 5 mice; Kv3.3KO = 6 from 3 mice and Kv3.1KO = 5 from 3 mice. **F:** AP amplitude, **G**: AP Decay slope, **H**: AP rise slope (10-90%) and **I**: membrane resistance for calyceal recordings. Average data presented as mean ± SD. Statistical test (parts E-I) were one-way ANOVAs and Tukey’s post hoc for multiple comparisons, with significant Ps indicated on the graph.

AP duration increased in terminals from Kv3.3KO mice, similar to that from WT mice in the presence of TEA. In contrast, the AP duration recorded from terminals of Kv3.1KOs was comparable to WT APs (see Figs. 1D & 1E). AP half-widths increased from 0.28±0.02 ms in WT (n=9) and 0.32±0.02 ms Kv3.1KOs (n=5) to 0.43±0.03 ms in Kv3.3KO mice (n=6) and 0.51±0.1 ms in WT+TEA (n=7; one-way ANOVA, Tukey’s post hoc; Kv3.3KO vs WT P=0.0001; Kv3.3KO vs Kv3.1KO P= 0.018; 1mM TEA vs WT P=0.0001). This increase in duration was accompanied by a slowed rate of AP decay in both Kv3.3KO terminals and WT terminals upon perfusion of TEA (Fig. 1G, one-way ANOVA, Tukey’s post hoc, Kv3.3KO vs WT P=0.0042; TEA vs WT P=0.0016) with no changes in the rising phase of the AP (one-way ANOVA, P=0.50; Fig. 1H), nor in the resting (input) membrane conductance (one-way ANOVA, P=0.56; Fig. 1I) or AP threshold (one-way ANOVA, P=0.96; see statistics table). This suggests that presynaptic Kv3.3 subunits are important for fast AP repolarization and presynaptic Kv3 channel formation. Despite the lack of a significant increase in action potential duration measured in Kv3.1KOs, the rate of decay was significantly slowed from 99±20 mV/ms (n=10) to 66±16 mV/ms (n=5; one-way ANOVA, Tukey’s post hoc, P=0.02), consistent with Kv3.1 having a secondary role in presynaptic AP repolarization, as would be expected from their localization at axonal nodes of Ranvier (Devaux ***et al.***, 2003).

### Deletion of Kv3.3 subunits increases calyceal-evoked EPSC amplitude

Transmitter release crucially depends on depolarization of the presynaptic membrane and consequent calcium influx. Since Kv3 channels are present and involved in calyceal AP repolarization, we assessed the physiological impact of each Kv3 subunit on transmitter release, and compared this to WT and to transmitter release following pharmacological block of presynaptic Kv3 channels using 1 mM TEA.

Whole cell patch recordings under voltage-clamp (HP = −40 mV, to inactivate voltage-gated Na^+^ channels) were made from MNTB neurons possessing an intact calyx, in tissue from mice of each genotype and in the presence of 1 mM TEA. This allows comparison of the EPSC amplitude in four conditions: where Kv3 channels are intact (WT) and where they lack Kv3.1 or Kv3.3, and finally with all presynaptic Kv3 channels blocked (TEA). Unitary calyceal EPSCs were evoked in MNTB principal neurons as shown in Fig. 2A & 2F. The global average for each condition and the unitary mean evoked EPSCs are overlaid from each genotype in Fig. 2F. The dashed grey line shows the WT mean amplitude for comparison. The calyceal EPSCs recorded from Kv3.3KO mice were of larger peak amplitude and longer lasting compared to WT or Kv3.1KOs, consistent with increased transmitter release from the calyx of Held in the Kv3.3KO.

**Figure 2:**
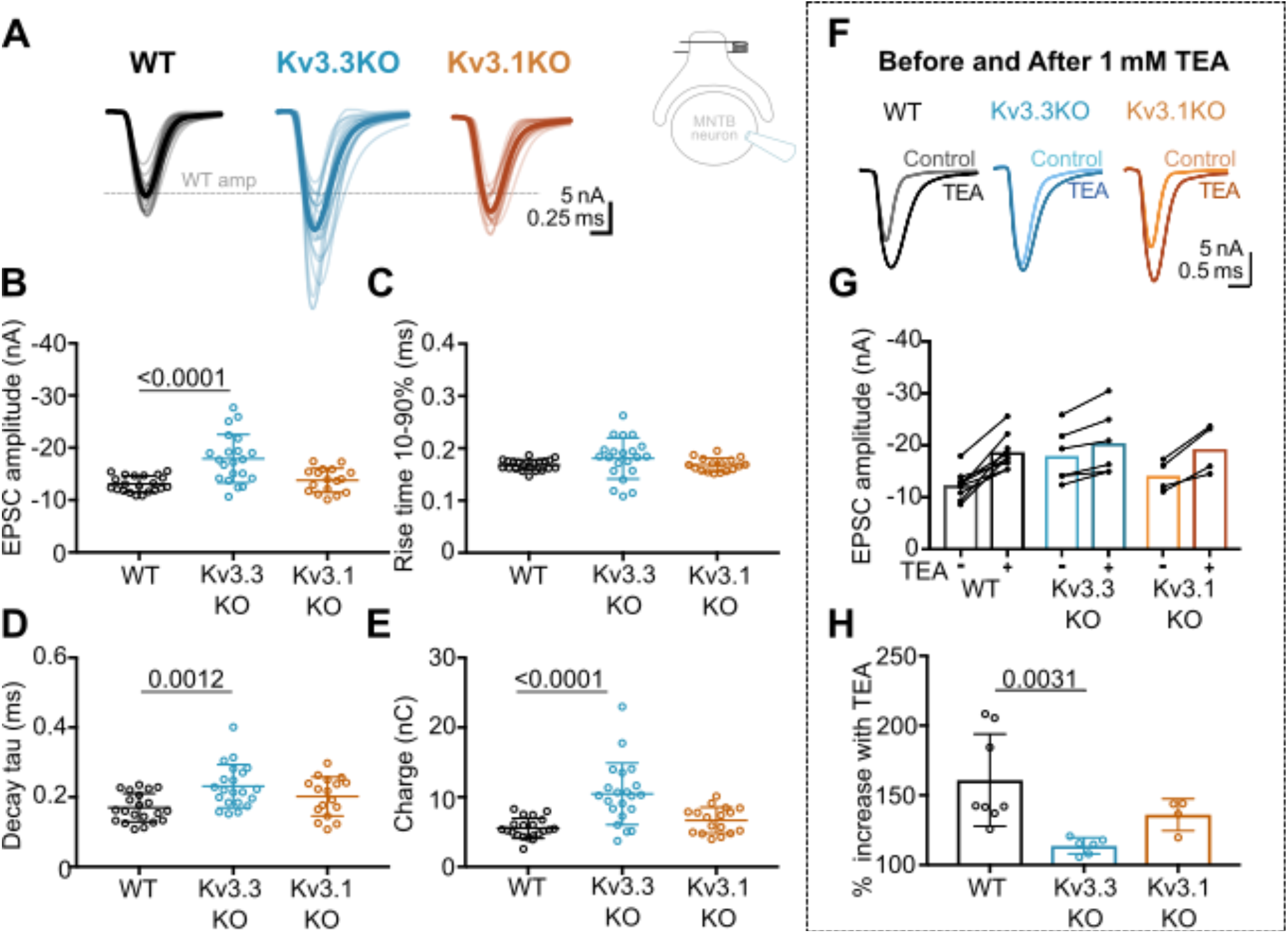
Kv3.3 deletion increases EPSC_s_ and occludes block by TEA. **A:** Superimposed calyceal EPSCs generated from each genotype (age P21-P25): wildtype (WT; black), Kv3.3KO (blue) and Kv3.1KO mice (orange). Thin lines show traces from individual neurons (each is mean of 5 EPSCs) with thick line showing the population mean for each genotype. Grey dashed line indicates the average WT amplitude; N = WT, 22 neurons (11 mice); Kv3.3KO, 22 neurons (10 mice); Kv3.1KO, 17 neurons (8 mice). Inset shows recording and stimulation configuration. **B:** EPSC amplitude, 10-90% rise time, decay tau and total charge of postsynaptic events. Significant increases were detected in amplitude, decay tau and charge of EPSCs from Kv3.3KO animals (P values reported on graphs). **C.** No differences were seen in rise time (10-90%) between groups (one-way ANOVA, P= 0.1576). **D:** EPSC decay tau and **E:** EPSC total charge were increased in the Kv3.3KO relative to WT. **F:** EPSC traces from WT, Kv3.3KO and Kv3.1KO mice, before and after the addition of 1 mM TEA. (**Centre):** EPSC amplitudes recorded before and after perfusion of TEA (1mM); n = WT, 9 neurons (7 mice); Kv3.3KO, 6 neurons (3 mice); Kv3.1KO, 5 neurons (3 mice). **G:** Increase in EPSC amplitude by 1 mM TEA. **H**. The amplitude increase induced by TEA was significantly reduced in Kv3.3KO mice compared to WT. Average data presented as mean ± SD; statistics was using one-way ANOVA with Tukey‘s post hoc for multiple comparisons. Kruskal-Wallis ANOVA with Dunn’s multiple corrections was used to compare change to EPSC amplitude in TEA due to a non-gaussian distribution in WT.

The data for EPSC peak amplitude, rise-time, decay tau and charge are plotted in Fig. 2B–2E, respectively, for each genotype. EPSC amplitudes were similar in WT and Kv3.1KO, - 13±2 nA (n=21) and −14±2 nA (n=17), respectively; but increased significantly in the Kv3.3KO to −18±5 nA (n=21; one-way ANOVA, Tukey‘s post hoc, Kv3.3KO vs WT, P= 0.0001; Kv3.3KO vs Kv3.1KO P=0.0005). The decay tau for the EPSCs in the Kv3.3KO slowed to 0.23±0.06 ms (n=21) compared to 0.17±0.04 ms in WT (n=21; one-way ANOVA, Tukey’s post hoc, P=0.0012). The total charge of EPSCs in Kv3.3KOs increased to −10±4 nC (n=21) compared to −6±1 nC and −7±2 nC in WT (n=21) and Kv3.1KOs (n=17), respectively (one-way ANOVA, Tukey’s post hoc; Kv3.3KO vs WT, P=0.0001; Kv3.3KO vs Kv3.1KO P=0.0007). No change of EPSC rise time was observed in either KO (one-way ANOVA, P=0.1918).

The increased EPSC amplitude observed in Kv3.3KOs suggests that Kv3.3 subunits are major contributors to repolarization of the presynaptic terminal. If Kv3.3 subunits dominate presynaptic Kv3 channels, then blocking presynaptic Kv3 channels with TEA will have a lesser effect on EPSCs from the Kv3.3KO. To test this hypothesis, we compared the effect of 1 mM TEA on EPSC amplitude from each of the three genotypes (Fig 3F–3H). Indeed, blocking Kv3 channels with TEA had a much smaller effect on EPSC amplitude in the Kv3.3 KO compared to WT and Kv3.1 KO. In WT, TEA increased EPSC amplitude by 160±33% (n=8), compared to 114±6% (n=6) in Kv3.3KO (Kruskal-Wallis, Dunn‘s multiple comparison, P=0.0031). The EPSC amplitude in Kv3.1KO (136±11%, n=4) was not significantly different from WT (Kruskal-Wallis, Dunn’s multiple comparison P=0.99). This result is consistent with dominance of presynaptic repolarization by channels containing Kv3.3.

**Figure 3:**
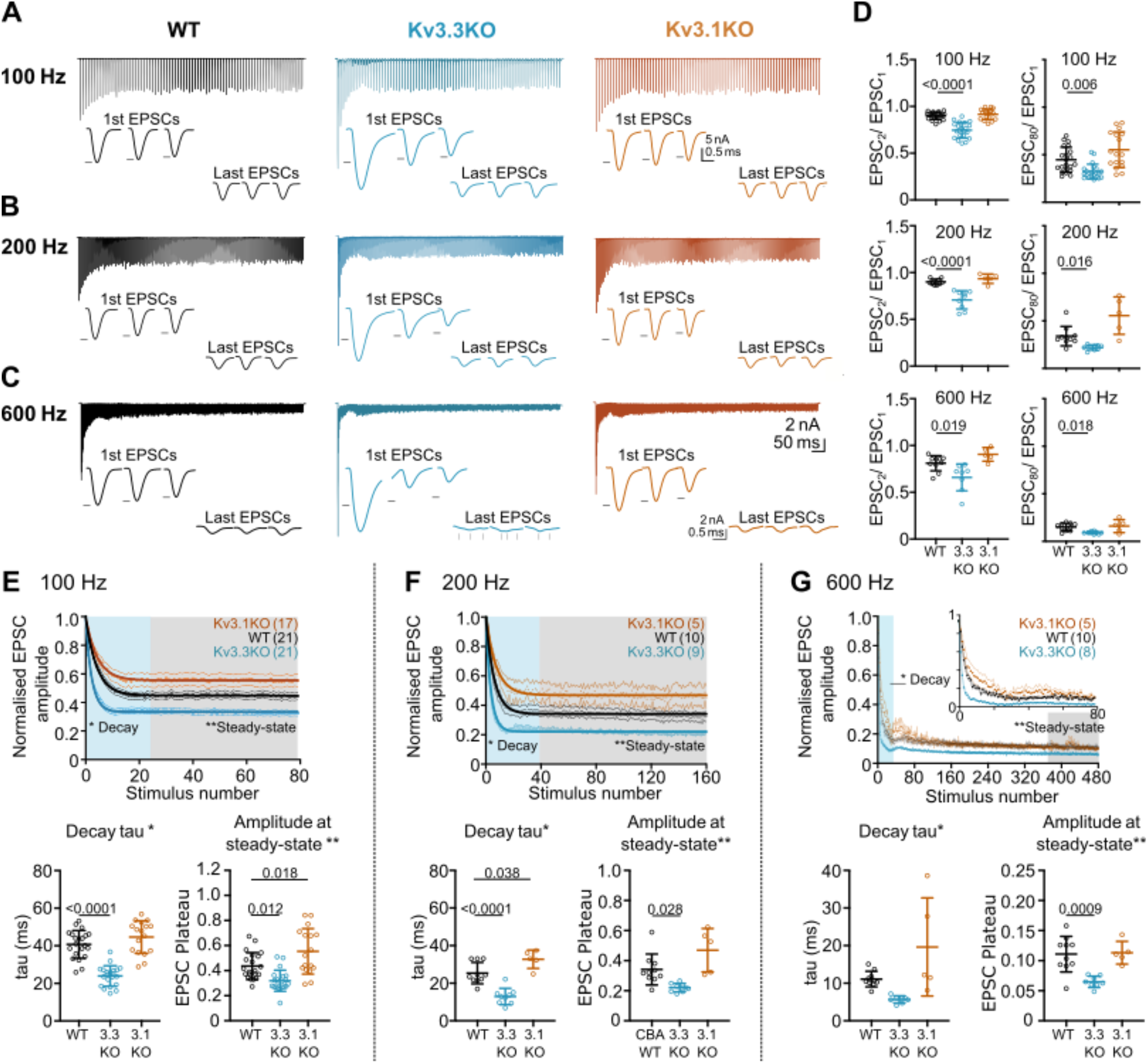
Evoked EPSC short-term depression is faster and more pronounced in the Kv3.3KO. **A,B,C:** Example MNTB EPSC trains (800 ms) on stimulation at **A:** 100, **B:** 200 or **C:** 600Hz for WT (left, black), Kv3.3KO (middle, blue) and Kv3.1KO (right, orange) mice (aged P21-P25). Each trace shows an average of 5 trials from a single neuron with stimulus artefacts removed for clarity. Lower insets: The first and last three EPSC traces are shown below each train. Black arrows show EPSC amplitudes from the WT mouse (left). Vertical grey arrows indicate multiple, asynchronous responses in the final EPSCs of Kv3.3KO traces. **D:** Paired-pulse ratios for EPSC2/EPSC1 (left) and EPSC80/EPSC1 (right) generated in response to 100 Hz (top), 200 Hz (middle) and 600 Hz (bottom); in each case the ratio is significantly decreased in Kv3.3KO with respect to WT. For 100Hz - WT: N=21cells (11 mice); Kv3.3KO: N=21 (10 mice); Kv3.1KO: N=17(8 mice); for 200 & 600 Hz - WT: n=10(6 mice); Kv3.3KO: n=9 (4 mice); Kv3.1KO: n=5 (3 mice). Data shows mean ± SD and statistical significance one-way ANOVAs with Tukey‘s post hoc for multiple comparisons or Kruskal-Wallis with Dunn’s multiple comparison (EPSC80/EPSC1 at 100 & 200 Hz due to non-gaussian data distributions), significant P values given on the respective graphs. **E, F, G:** Normalized EPSCs, short-term depression is faster and larger in Kv3.3KO mice (compared to WT and Kv3.1KO mice) **E:** 100 Hz **F:** 200 Hz **G:** 600 Hz. The rate of EPSC depression is plotted as a single exponential tau (lower left) and normalized EPSC amplitudes at steady-state are plotted (lower right) for each genotype and stimulus frequency. N numbers are presented in brackets on normalized EPSC amplitude graphs (and are the same neurons as used in D). One-way ANOVAs with Tukey’s post hoc for multiple comparison correction were used to test significance, P values reported on graphs. All data plotted as mean ± SD, except in graphs of normalized EPSC amplitudes (E, F and G, top) where data is plotted as mean ± SEM for clarity.

### Enhanced short-term depression in Kv3.3KO

The calyx of Held/MNTB synapse is capable of sustained transmission at firing frequencies of around 300Hz *in vivo* (Kopp-Scheinpflug ***et al.***, 2008) (also see Fig. 7), encoding information about sound stimuli for binaural integration with high temporal precision. Indeed, the calyx of Held giant synapse can sustain short AP bursts with peak firing rates of up to 800Hz, *in vitro* (Taschenberger & Gersdorff, 2000) for a few milliseconds. Clearly, the increased transmitter release observed in the Kv3.3KO (Fig. 3) has consequences for maintenance of EPSC amplitude during repetitive firing, in that vesicle recycling and priming must rapidly replace docked vesicles if transmitter release is to be maintained for the duration of the high frequency train. Repetitive stimulation shows that the evoked EPSC amplitude declines during a stimulus train until rates of vesicle priming are in equilibrium with the rate of transmitter release for a given stimulus frequency. To assess repetitive transmitter release in each of the genotypes we evoked EPSCs over a range of frequencies from 100 to 600 Hz (with each stimulus train lasting 800 ms).

Whole cell patch recordings from voltage-clamped MNTB neurons (HP = −40mV) were conducted in which calyceal EPSC trains were evoked at 100 Hz, 200 Hz or 600 Hz. Each train was repeated three times, with a resting interval of 30s between repetitions of stimuli trains. The EPSC amplitudes during the trains were compared in MNTB neurons of each genotype: WT (black), Kv3.3KO (blue) and Kv3.1KO (orange) mice. The first EPSC amplitude in trains delivered to the calyx/MNTB from the Kv3.3KO mouse was of larger amplitude than observed in WT or in the Kv3.1KO and subsequent EPSCs showed a larger short-term depression. In Fig. 3A - 3C a matrix of EPSC trains are plotted with the same stimulus frequency in each row for each genotype. The inset traces show the first 3 EPSCs and the last 3 EPSCs in each train, with the black arrow heads indicating the WT amplitude for comparison across genotypes.

Paired pulse ratio (PPR) was significantly increased in the Kv3.3KO compared to both WT and Kv3.1KO at 100 Hz, 200Hz and 600Hz, with mean PPR (EPSC2/EPSC1) plotted in Fig 3D (left column). This enhanced depression of EPSC amplitude was maintained for the duration of the train shown in comparison with the 80^th^ EPSC in the train (EPSC80/EPSC1; Fig 3D, right column; Table 1).

**Table 1:**
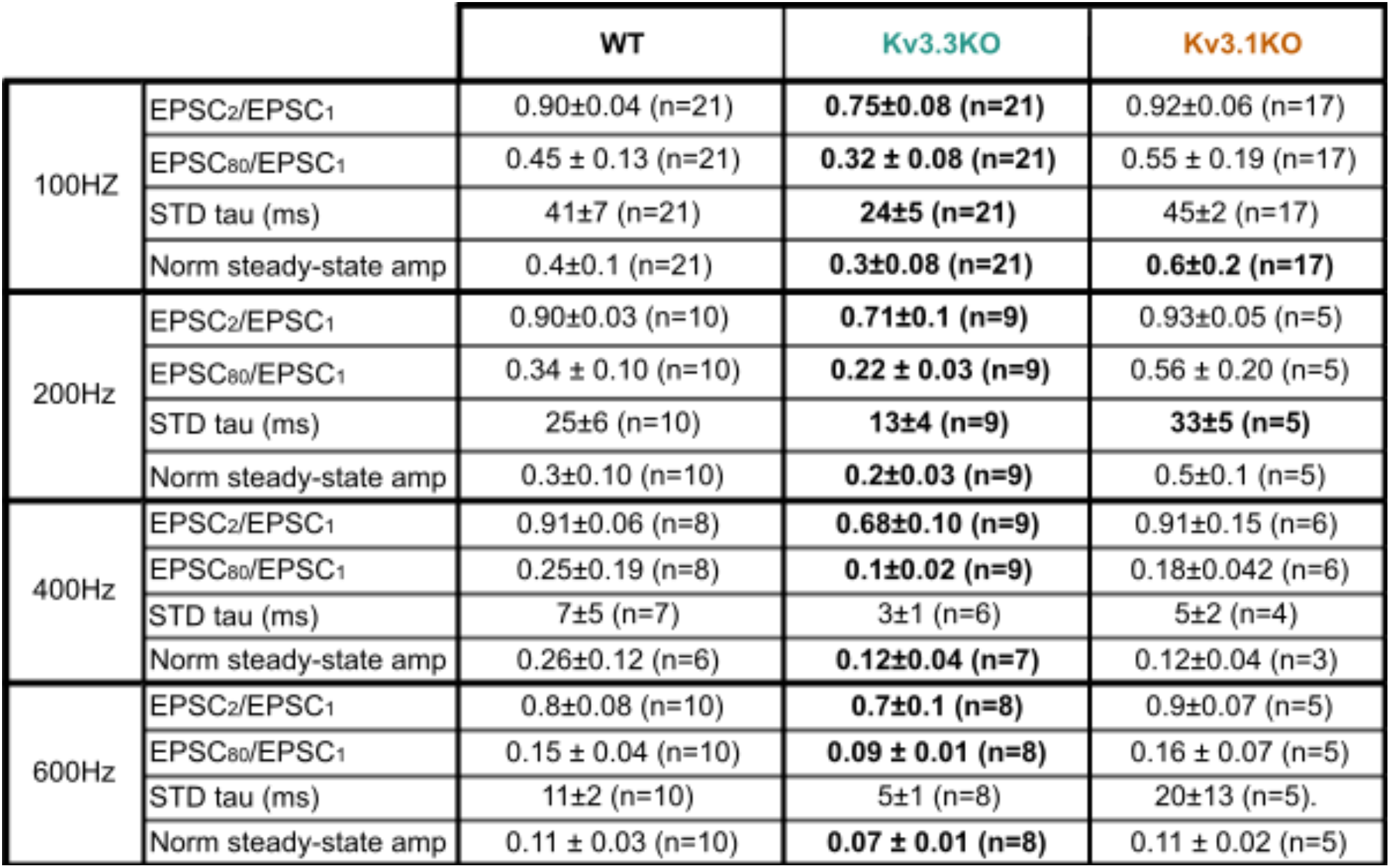
Short-term depression is increased and accelerated in mice lacking Kv3.3. Values shown for parameters measured in figure 3 for WT, Kv3.3KO and Kv3.1KO at 100, 200, 400 and 600Hz. Paired pulse depression of EPSC responses recorded in MNTB neurons (EPSC_2_/EPSC_1_) was increased in Kv3.3 KO animals during high frequency stimulation of calyx fibers. The increased depression was maintained throughout the stimulation train (EPSC_80_/EPSC_1_) across all frequencies. The rate of short term-depression in EPSC amplitudes during EPSC trains (800ms), measured as short-term depression (STD) decay tau was significantly increased in Kv3.3 KOs at 100 and 200Hz compared to WT. This STD was more severe in mice lacking Kv3.3, shown by decreased normalized steady-state EPSC amplitudes compared to WT. STD tau and steady state amplitudes were measured using a single exponential fit to normalized EPSC amplitudes throughout the 800ms stimulation trains. N=number of neurons. Values in bold are significantly different to WT (see statistics table for more detail). Data represented as mean ± SD.

|  |  | WT | Kv3.3KO | Kv3.1KO |
| --- | --- | --- | --- | --- |
| 100Hz | EPSC<sub>2</sub>/EPSC<sub>1</sub> | 0.90±0.04 (n=21) | **0.75±0.08 (n=21)** | 0.92±0.06 (n=17) |
|  | EPSC<sub>80</sub>/EPSC<sub>1</sub> | 0.45 ± 0.13 (n=21) | **0.32 ± 0.08 (n=21)** | 0.55 ± 0.19 (n=17) |
|  | STD tau (ms) | 41±7 (n=21) | **24±5 (n=21)** | 45±2 (n=17) |
|  | Norm steady-state amp | 0.4±0.1 (n=21) | **0.3±0.08 (n=21)** | **0.6±0.2 (n=17)** |
| 200Hz | EPSC<sub>2</sub>/EPSC<sub>1</sub> | 0.90±0.03 (n=10) | **0.71±0.1 (n=9)** | 0.93±0.05 (n=5) |
|  | EPSC<sub>80</sub>/EPSC<sub>1</sub> | 0.34 ± 0.10 (n=10) | **0.22 ± 0.03 (n=9)** | 0.56 ± 0.20 (n=5) |
|  | STD tau (ms) | 25±6 (n=10) | **13±4 (n=9)** | **33±5 (n=5)** |
|  | Norm steady-state amp | 0.3±0.10 (n=10) | **0.2±0.03 (n=9)** | 0.5±0.1 (n=5) |
| 400Hz | EPSC<sub>2</sub>/EPSC<sub>1</sub> | 0.91±0.06 (n=8) | **0.68±0.10 (n=9)** | 0.91±0.15 (n=6) |
|  | EPSC<sub>80</sub>/EPSC<sub>1</sub> | 0.25±0.19 (n=8) | **0.1±0.02 (n=9)** | 0.18±0.042 (n=6) |
|  | STD tau (ms) | 7±5 (n=7) | 3±1 (n=6) | 5±2 (n=4) |
|  | Norm steady-state amp | 0.26±0.12 (n=6) | **0.12±0.04 (n=7)** | 0.12±0.04 (n=3) |
| 600Hz | EPSC<sub>2</sub>/EPSC<sub>1</sub> | 0.8±0.08 (n=10) | **0.7±0.1 (n=8)** | 0.9±0.07 (n=5) |
|  | EPSC<sub>80</sub>/EPSC<sub>1</sub> | 0.15 ± 0.04 (n=10) | **0.09 ± 0.01 (n=8)** | 0.16 ± 0.07 (n=5) |
|  | STD tau (ms) | 11±2 (n=10) | 5±1 (n=8) | 20±13 (n=5). |
|  | Norm steady-state amp | 0.11 ± 0.03 (n=10) | **0.07 ± 0.01 (n=8)** | 0.11 ± 0.02 (n=5) |

The mean data, normalized to the amplitude of the first EPSC, is plotted for each genotype at the stated stimulus frequency: Fig. 3E–G (100 Hz, to 600 Hz). There are two key parameters plotted below each depression curve: first, the short-term depression decay time-constant (Decay Tau, defined as the rate at which the EPSC amplitude equilibrates to the new steady-state amplitude at each stimulus frequency); and second, amplitude at steady-state (the amplitude of the new steady-state EPSC during the train, relative to EPSC1). The Kv3.3KO consistently showed the fastest rate of short-term depression, compared to WT and Kv3.1KOs; this was highly significant at 100 Hz and 200Hz (Table 1).

This trend continued so that during stimuli at 600 Hz (Fig. 3G), a further increase in the rate of short-term depression was observed; again, this was most marked in EPSC trains from the Kv3.3KO compared to both WT and Kv3.1KO. The decay time-constant for short-term depression was 6±1 ms for Kv3.3KOs (n=8), 11±2 ms for WT (n=10) and 20±13 ms for Kv3.1KOs (n=5; Table 1).

Following short-term depression, the ‘steady-state’ EPSC amplitude was achieved after 20-40 evoked responses (Fig 3E, 3F, 3G; lower right graph). This reflects the net equilibrium achieved under control of four key presynaptic parameters: probability of transmitter release, the rate of AP stimulation, the rate of vesicle replenishment at the release sites and the size of the vesicle pool (see modelling section below). In young animals, postsynaptic AMPAR desensitization can also play a role in short-term depression, but this is a minor contribution at mature synapses and physiological temperatures, as employed here (Taschenberger ***et al.***, 2002; Wong ***et al.***, 2003). At all frequencies, EPSCs from the Kv3.3KO mice showed lower amplitude steady-state values in comparison to WT; while the Kv3.1KO data was either the same or greater than WT (Fig. 3E, Table 1).

### Kv3.3 deletion accelerated a fast component of recovery from short-term depression

Presynaptic spike broadening has previously been shown to enhance vesicle recycling during repetitive stimulation trains (Wang and Kaczmarek, 1998) due to enhanced calcium dependent recovery. An increase in the rate of recovery of the evoked EPSC following short-term depression (Fig 4.) could indicate changes in vesicle recycling. In WT animals the EPSC was depressed to around 40% on stimulation at 100Hz (Fig. 3E), increasing to 90% depression for 600Hz (Fig. 3G). Three example conditioning traces are shown in Fig. 4A–C for each genotype. On ceasing stimulation, the depressed EPSC recovered back to control amplitudes over a time-course of 30 seconds (Fig. 4A–C, recovery). The recovery phase was measured by presenting stimuli at intervals of: 50, 100, 200, 500 ms, and 1, 2, 5, 10, 20 and 30 s after the end of the conditioning train. Each recovery curve was repeated 3 times and the mean EPSC amplitudes plotted over the 30 second period (Fig. 4D) and fit with a double exponential. The fast time-constant contributed around half of the recovery amplitude and this did not differ significantly between the three genotypes (Fig. 4E). The fast time-constant was significantly accelerated in the Kv3.3 KO compared to WT and Kv3.1 KO (0.2 ±0.2 s in Kv3.3KO; 0.45 ± 0.2 s in WT and 0.4 ±0.2 s in the Kv3.1KO; Fig. 4F, p = 0.025). The slow time-constant was 5±3 s in WT, and was similar to both Kv3.1KO and Kv3.3KO (Fig. 4G). The enhanced fast recovery rate is consistent with an activity-dependent component of vesicle recycling as observed previously on blocking presynaptic Kv3 channels at the calyx of Held (Wang & Kaczmarek, 1998).

**Figure 4:**
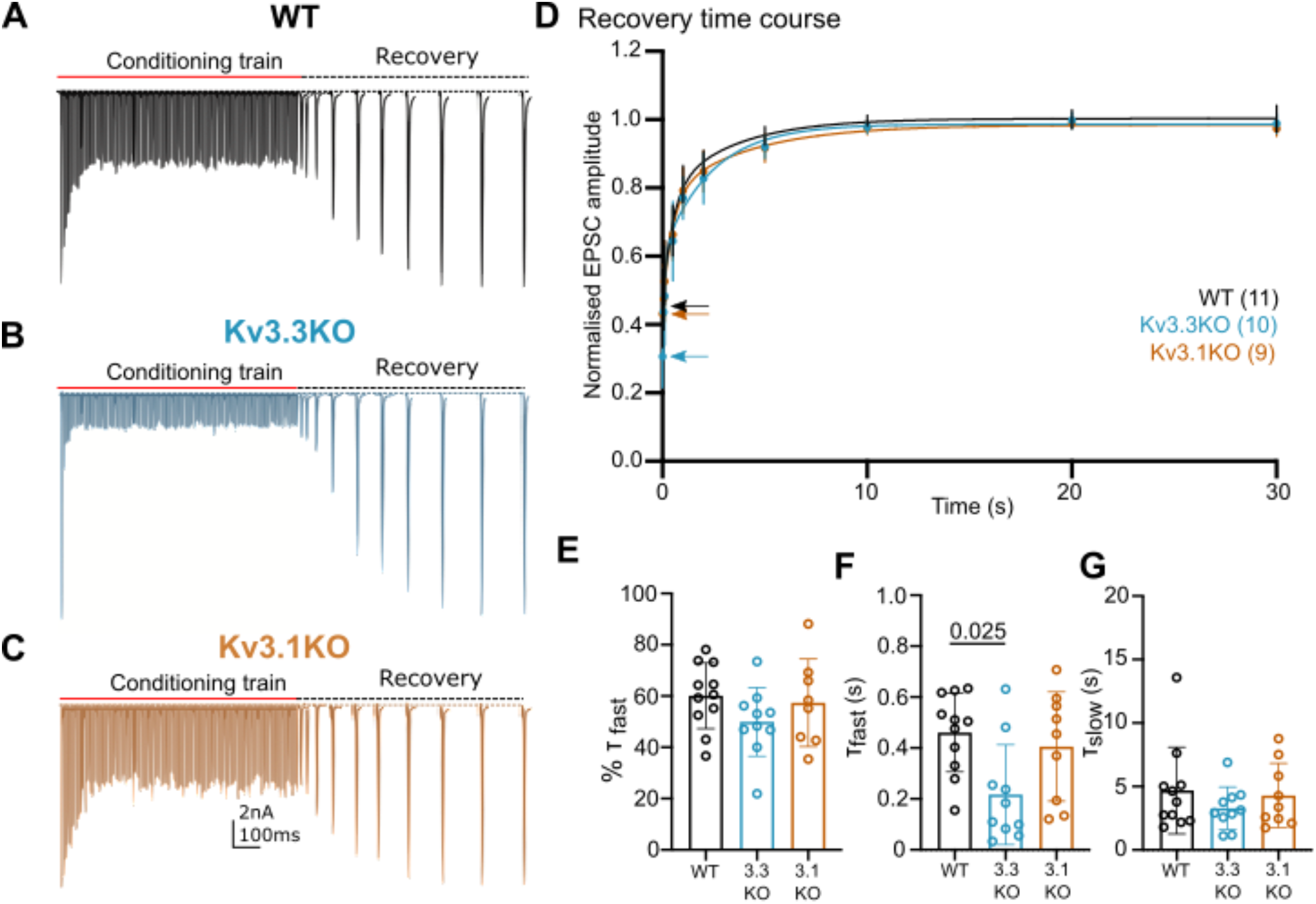
Recovery from short-term depression is accelerated on deletion of Kv3.3. **A:** WT (black), **B:** Kv3.3KO (blue), **C:** Kv3.1KO (orange). A representative example is shown for each genotype. A conditioning train of 100Hz (800 ms duration) evoked EPSCs displaying short-term depression. The recovery was estimated by delivery of single stimuli at intervals following the conditioning train (50ms, 100ms, 500ms, 1s, 2s, 5s, 10s, 20s, and 30s. Recovery intervals not to scale). **D:** The mean EPSC amplitude during the recovery is plotted for each genotype (mean ± SD. WT, black; Kv3.3KO, blue; Kv3.1KO, orange) over the 30s recovery period. The mean amplitude at the end of the conditioning train, from which the recovery starts, is shown by the respective coloured arrow. A double exponential was fit to each individual recovery curve and the mean curve is plotted for the respective genotype. Values are plotted as raw data and mean ± SD in E-G; N values from part D also apply here. **E:** The percent contribution of the fast component is similar between genotypes. **F:** The fast time-constant significantly accelerated from 0.4s in WT to 0.2s in the Kv3.3KO. **G:** The slow time-constant at around 5 seconds was unchanged between genotypes.

### Computational Model of transmitter release

The magnitudes and rates of EPSC depression and recovery following synaptic train stimulation provided constraints in refining a computational model of transmitter release and vesicle recycling at the calyx of Held (Graham ***et al.***, 2004). A simple model of transmission at the calyx of Held, based on transmitter release and recycling was employed as set out in Fig. 5A, equations 1–3. It included activity-dependent vesicle recycling (Graham ***et al.***, 2004; Billups ***et al.***, 2005; Lucas ***et al.***, 2018) and parameters were fit across the range of stimulus frequencies (100-600Hz) for the WT and the Kv3.3KO mouse. The model possessed a readily releasable pool (RRP; normalized size n), from which vesicles undergo evoked exocytosis with a release probability (Pv), following each AP and are recycled with a time-constant τ_r_. A basal rate of recycling (large time-constant τ_b_) is accelerated to an activity-dependent rate (smaller time-constant τ_h_) on invasion of an AP, and relaxed back to τ_b_ with a time-constant of τ_d_. The table in Fig. 5B shows that the dominant change in the model parameters between WT and Kv3.3KO, was an increase in the probability of vesicular release (Pv) from 0.13 in WT to 0.266 in the Kv3.3KO. There was also evidence for an increase in activity-dependent recycling, with a 20% acceleration of the replenishment rate (smaller τ_h_ in Kv3.3KO). The fit of the model to the mean experimental data (±SEM) is shown for the short-term depression and the recovery curves (WT: Fig. 5C–5D; Kv3.3KO: Fig. 5E–5F). This model showed that the physiologically observed changes in short-term depression and recycling could be fit across the range of stimuli rates with only two parameter changes in the Kv3.3KO: a dramatically increased Pv and a modest decrease in the activity-dependent vesicle recycling time-constant τ_h_ (higher recycling rate).

**Figure 5:**
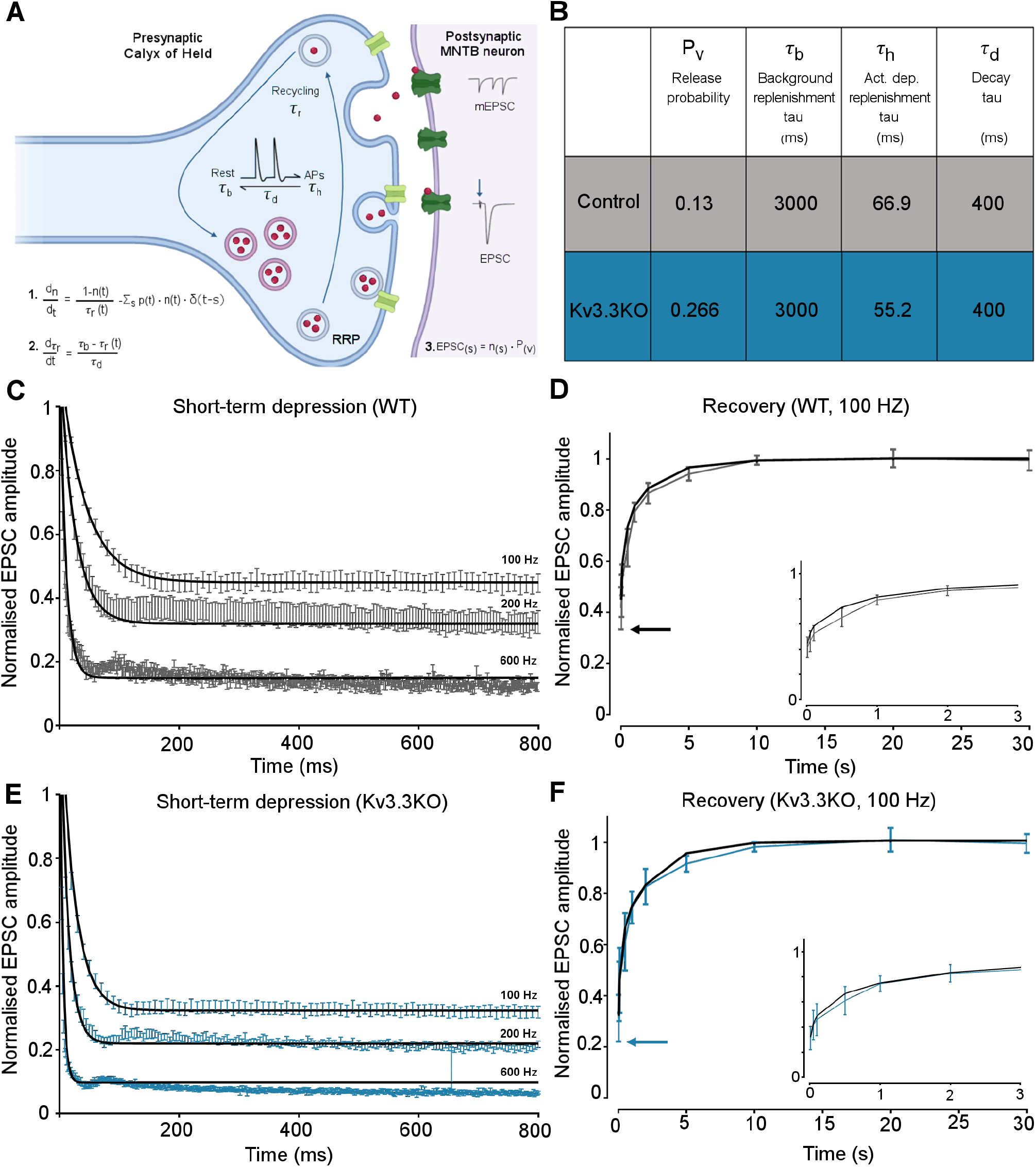
Kv3.3 deletion increases release probability and speeds a fast component of vesicle recycling, based on computational modelling. **A**. Model illustration: Vesicles are released from the readily releasable pool (RRP; normalized size n) with a probability Pv; the RRP is replenished with the recycling time-constant τ_r_. In the absence of APs τ_r_ = τ_b_ – the background replenishment time-constant, which decreases (accelerates) to τ_h_ following a presynaptic AP, and decays back to τ_b_ with a time-constant τ_d_. Model parameters were fit to WT data for evoked EPSC trains (100-600Hz) and their 100Hz recovery curves. **B.** Table showing the model parameters for fitting WT and Kv3.3KO data. Increasing Pv from 0.13 to 0.266 and accelerating τ_h_ from 66.9 to 52.2 were sufficient to fit the changes observed on Kv3.3 deletion. **C.** WT: EPSC amplitude data (mean ± SEM) during the conditioning train (100, 200 & 600Hz, grey) are plotted with superimposed model prediction curves (black). **D**. WT 100 Hz: Recovery of the EPSC (mean ± SEM) over 30 seconds. Inset shows data and fit for the first 3 seconds. Model fit is the superimposed black line. Horizontal arrow indicates EPSC amplitude at the end of the conditioning train. **E.** Kv3.3KO: EPSC amplitude data (mean ± SEM) during the conditioning trains are plotted (blue) with superimposed model prediction curves (black). **F**. Kv3.3KO 100 Hz: Recovery of the EPSC amplitude (mean ± SEM) over 30 seconds. Inset shows data and fit for the first 3 seconds. Model fit is the superimposed line. Horizontal arrow indicates EPSC amplitude at the end of the conditioning train.

### Integration of EPSCs in generating APs in the postsynaptic MNTB neuron

A key physiological question is the extent to which presynaptic Kv3.3 influences AP firing of the MNTB neuron in response to trains of synaptic stimuli. The initial observation is that MNTB AP output in response to high frequency calyceal EPSC input, declines with both Kv3.1 and Kv3.3 deletion (Fig. 6A) when measured over the whole train (WT: 52.49±8.15%, n=5; Kv3.3KO: 29.8±10.43%, n=7; Kv3.1KO: 36.29±11.12%, n=6; Kv3.3KO vs WT P=0.0044, Kv3.1KO vs WT P=0.046, one-way ANOVA, Tukey’s post hoc). But closer inspection reveals three phases of AP firing to evoked trains of calyceal synaptic responses as illustrated in Fig. 6B; which shows MNTB EPSCs/APs during an 800ms 600Hz train in a WT mouse. In phase I (green trace, Fig.6B) every evoked EPSP triggered one MNTB AP, so the MNTB output matched the calyx input. Firing then transitioned to Phase II (blue trace, Fig. 6B) after around 6-9 stimuli, where EPSPs often failed to evoke an AP, and the MNTB firing becomes chaotic and unpredictable. In Phase III (black trace, Fig. 6B): the EPSP reached a ‘steady-state’ amplitude, and the MNTB neuron fired APs to alternate EPSPs. The duration of Phase I was essentially identical across the three genotypes (Fig. 6C) at frequencies up to 200Hz, all showed 100% firing throughout the train; then from 300Hz and above, the Phase I duration declined dramatically for all genotypes. Genotype-specific limitations were observed at the highest frequencies. Fig. 6D shows the relative duration (%) of each phase for 600Hz EPSP trains. There were no differences in the duration of phase I, while Phase II and III were of variable duration. At 600Hz stimulation frequency the latency to the start of Phase II was 34.3±12.3ms for WT (n=10), 25±8 ms for Kv3.3KO (n=6) and 26±14ms for Kv3.1KO (n=5). The latency for Phase III was similar in WT and Kv3.1KO (510±224 ms and 597±251 ms, respectively) but in the Kv3.3KO only 1 out of 6 calyx/MNTB pairs briefly entered Phase III firing (Kv3.3KO vs WT Phase II P=0.0135, Phase III P=0.02, two-way ANOVA, Tukey’s post hoc). This is consistent with the idea that presynaptic Kv3.3 and hence fast APs, serve a role in maintaining information transmission across the synapse during high frequency firing, when short APs conserve resources, and reduce the rate of short-term depression.

**Figure 6:**
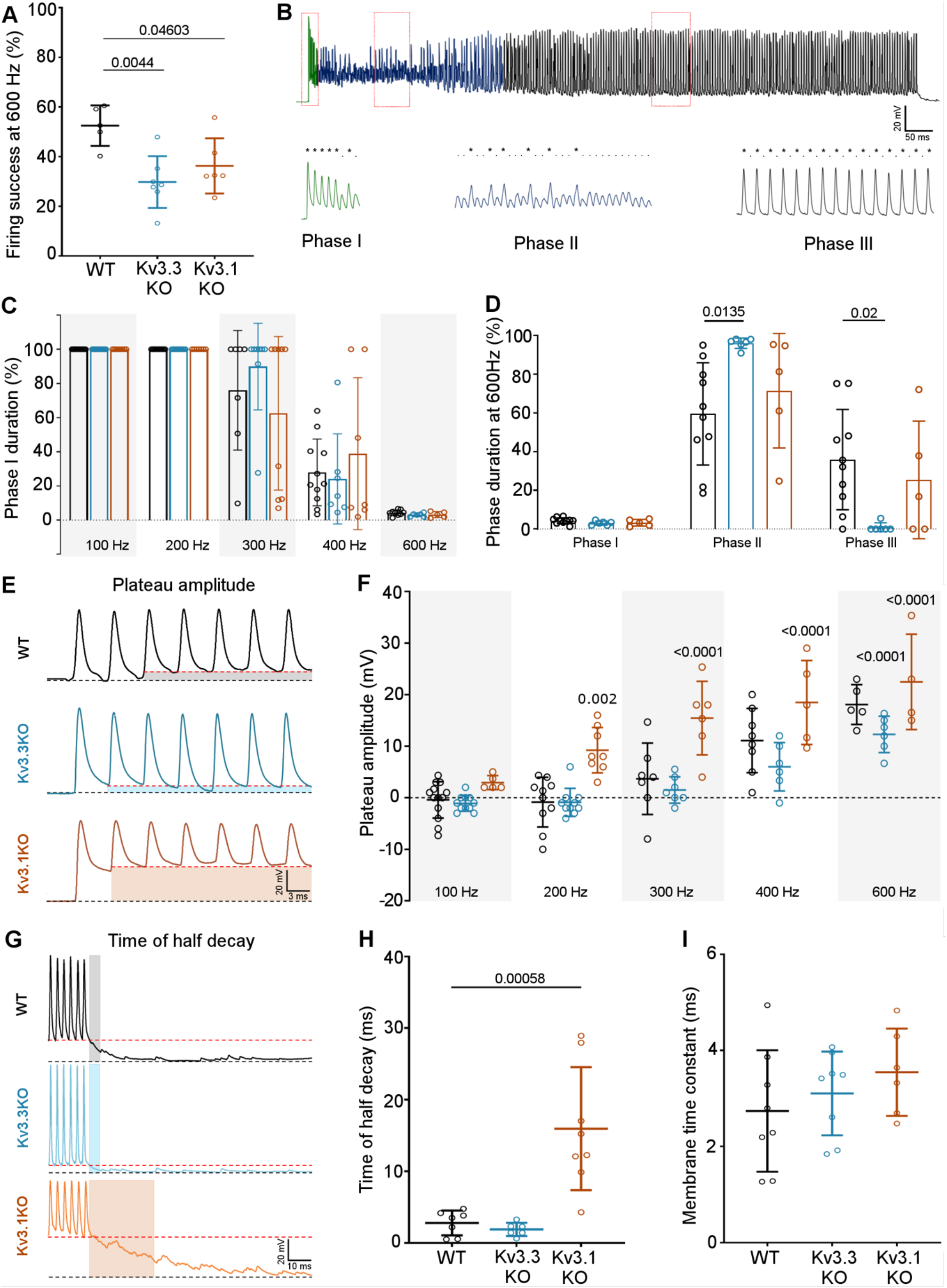
Kv3.3 deletion reduced ability to sustain MNTB AP firing at high frequencies. **A:** Percent firing success of MNTB neurons for 600Hz calyceal stimulation is reduced for both Kv3.3KO and Kv3.1KO. **B.** A representative AP train recorded for an MNTB neuron in response to 600 Hz synaptic stimulation lasting 800ms. Three phases of input:output firing defined: **Phase I** (green) 1:1 AP firing calyx:MNTB for each EPSC is prevalent early in the train. **Phase II** (dark blue) follows phase I where some EPSCs drop below threshold, and AP firing becomes less probable and chaotic. **Phase III** (black) MNTB AP firing becomes regular again but is now firing to alternate input EPSCs, restoring information transmission. Expanded sections (boxed) of each phase of AP firing are shown below the full trace, APs are indicated by ‘*‘ and EPSPs that are below threshold are indicated by ‘.’. **C:** Phase I duration across stimulation frequencies (100-600Hz) for each genotype (WT, black; Kv3.3KO, blue; Kv3.1KO, orange): Phase I firing lasts throughout the train at frequencies up to 300Hz, but declines to only a few milliseconds at 600Hz synaptic stimulation, but there were no significant differences between the genotypes. **D:** The time spent in each phase for 600Hz stimulation train (with each genotype indicated by the same colours as in C). The MNTB neuron is unable to maintain Phase I and Phase II dominates for each genotype. However, Phase III is only achieved briefly on 1 of 6 observations in the Kv3.3KO. **E:** A sustained depolarized plateau was also observed, in the MNTB AP trains, as indicated by the shaded regions in this data from 300Hz, and was particularly large the Kv3.1KO. **F:** The amplitude of the depolarization plateau increased in magnitude with stimulation frequency for each genotype, but was significantly larger in the Kv3.1KO at all frequencies above 100Hz. **G:** A slowly decaying depolarization following the end of the synaptic train, as shown for representative traces from each genotype. **H:** This decaying depolarization is quantified as the time to half decay and was significantly longer in the Kv3.1KO compared to WT in a 300Hz AP train. **I:** The postsynaptic MNTB neuron membrane time constant was unchanged across all genotypes (genotype: WT, black; Kv3.3KO, blue; Kv3.1KO, orange).

Large magnitude Kv3 currents in postsynaptic MNTB neurons demonstrably assists in transmission of timing information (Song ***et al.***, 2005). Kv3 has little impact on resting MNTB membrane time-constant or AP firing threshold (induced by current injection through the pipette) and these are essentially identical in the three genotypes (Choudhury ***et al.***, 2020), which was also confirmed here (Fig. 6i). However, in the Kv3.1KO, calyceal stimulation evoked a sustained depolarization (in addition to EPSPs) at frequencies above 100Hz (Fig. 6E). The mean amplitude of this depolarizing plateau potential during the train is plotted against stimulus frequency for each genotype (Fig. 6F). Although all genotypes exhibited this plateau depolarization at 600Hz, it was significantly larger for the Kv3.1KO at frequencies above 100Hz (Fig. 6F) In contrast, Kv3.3KOs showed a reduced plateau potential at frequencies above 300Hz, reaching significance only at 600Hz (Fig. 6F; two-way ANOVA, Tukey’s post hoc). This was also observed as a decaying depolarization at the end of the train (Fig. 6G), with the time to half-decay plotted in Fig. 6H (WT: 2.79±1.73 ms, n=7: Kv3.3KO: 1.89±0.92 ms, n=6: Kv3.1KO: 15.96±8.59 ms, n=8: Kv3.1KO vs WT P=0. 000553, one-way ANOVA, Tukey’s post hoc). In WT and Kv3.3KO genotypes, this decay was similar to the postsynaptic membrane time-constant, but in the Kv3.1KO, non-synchronous spontaneous EPSPs were observed. The membrane time constants for all genotypes were essentially identical, consistent with no change in glutamate receptor expression (WT: 2.74±1.26ms, n=8; Kv3.3KO: 3.1±0.87ms, n=8; Kv3.1KO: 3.54±0.9ms, n=6). This plateau depolarization caused increased depolarization block of APs and thereby undermined our ability to examine AP firing physiology in the Kv3.1KO. Therefore *in vivo* experiments focused on comparison of the WT and Kv3.3KO genotypes, which did not exhibit this postsynaptic epi-phenomenon.

### Kv3.3 increases temporal precision and signal-to-noise ratios in response to sound

The prolonged presynaptic AP duration and increased transmitter release observed here in the Kv3.3KO, shows that the calyx has a specific need for Kv3.3 channels. So what is the impact of presynaptic Kv3.3 on auditory processing?

Behaviourally, the Kv3.3KO mouse is as sensitive to sound as the WT in that Auditory Brainstem Response (ABR) thresholds were similar in 6 monthold WT and Kv3.3KO mice, with no statistical difference across a wide frequency range (see summary statistics, unpaired t-test with Holm-Šídák’s test for multiple comparisons). This provided the opportunity to examine auditory processing in the MNTB *in vivo* using extracellular recordings from WT and Kv3.3KO mice during sound stimulation. This data was non-gaussian and summary statistics are presented as median, and quartile values (in square brackets).

Extracellular MNTB single unit recordings exhibited a typical complex waveform, comprised of a presynaptic and a postsynaptic component (Fig. 7A) (Kopp-Scheinpflug ***et al.***, 2003). The time between the peak and trough of extracellular APs is a compelling marker for AP halfwidth (Ritzau-Jost ***et al.***, 2021) and confirmed our results of the presynaptic patch clamp recordings. AP halfwidth of the presynaptic AP (preAP) was significantly longer in Kv3.3KOs (0.25ms [0.16; 0.31]; n=13) compared to WT recordings (0.17ms [0.16; 0.19]; n=20; Fig. 7B; Mann-Whitney Rank Sum Test: *P*=0.036). Synaptic delays as measured by peak-to-peak times in the complex waveform were also significantly prolonged in Kv3.3KOs (0.56ms [0.45; 0.70]; n=13) compared to WT controls (0.44ms [0.41; 0.49]; n=20; Fig. 7C; n=13; Mann-Whitney Rank Sum Test: *P*=0.013). While changes in presynaptic AP duration and synaptic delay may predominantly affect temporal processing, the prolonged postsynaptic AP duration observed in the Kv3.3KOs (0.64ms [0.29; 0.46]; n=13; Fig.7D) might influence high-frequency firing abilities (WT: 0.36ms [0.48; 0.82]; n=20; Mann-Whitney Rank Sum Test: *P*≤0.001).

**Figure 7:**
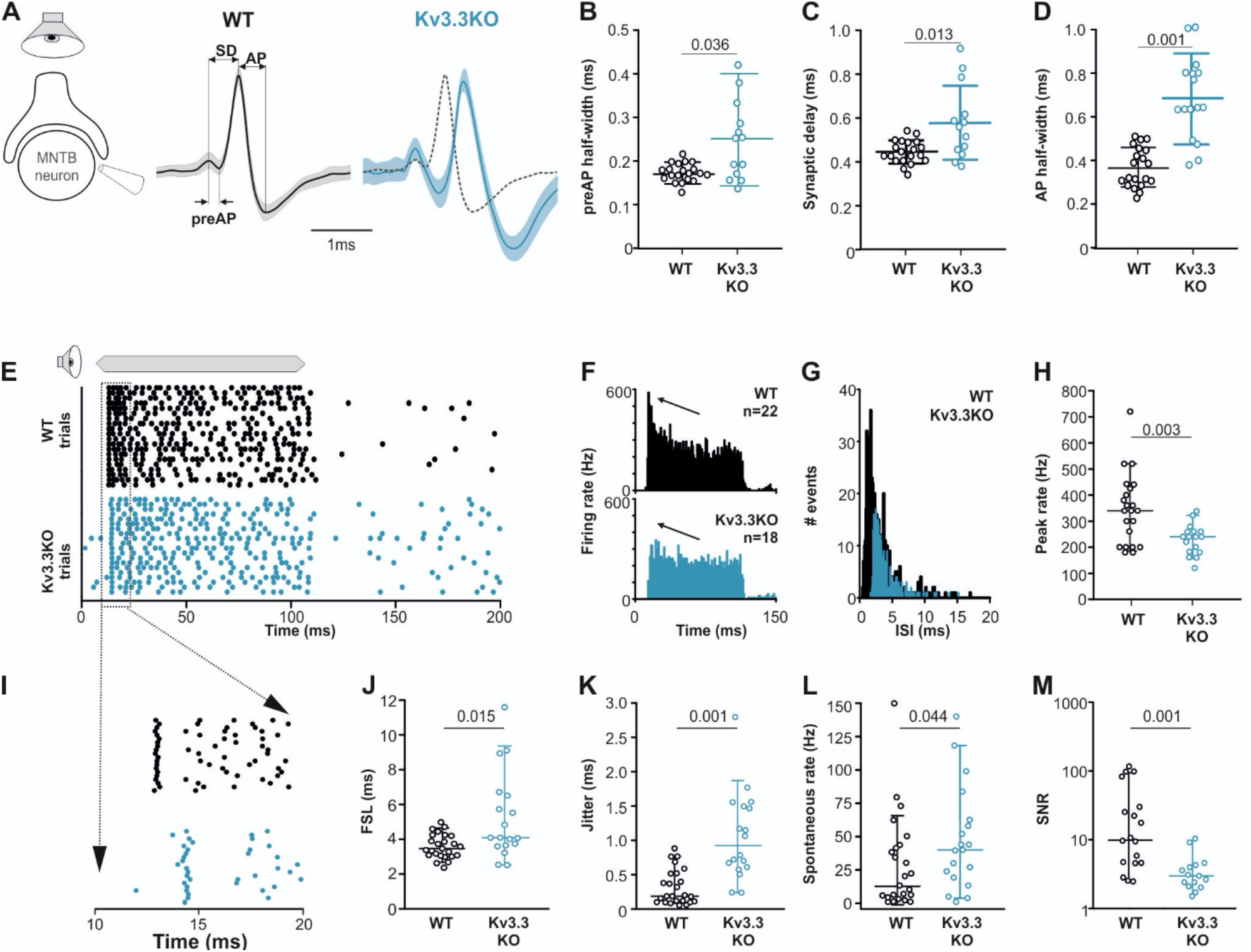
Presynaptic Kv3.3 accelerates the brainstem response to sound and improves timing and signal-to-noise ratio. **A.** Extracellular recording from calyx/MNTB *in vivo* shows complex APs (presynaptic and postsynaptic) from WT (black) and Kv3.3KO (blue) mice in response to sound; overlay of APs (right) shows delayed and longer APs in the KO. **B. C. D.** Presynaptic AP halfwidth, synaptic delay and postsynaptic AP halfwidth are all longer in the Kv3.3KO (blue) than WT (black). **E.** Raster display of MNTB AP response to sound (20 trials, 100ms duration) and spontaneous firing for both WT (black, upper) and Kv3.3KO (blue, lower). **F. G. H.** Peri Stimulus Time Histogram (PSTH) of the evoked APs (grand avg, 1ms bins) show reduced peak firing rates (F, black arrow) due to longer interspike intervals (ISIs) in the Kv3.3KO (blue) and the peak firing rate is significantly reduced. **I. J. K.** Expansion of first 20 ms of the Raster plot shows increased first spike latency and jitter (latency SD) in the Kv3.3KO (blue). **L. M.** The mean spontaneous firing rate was higher in the Kv3.3KO (blue) and overall, these changes degraded signal-to-noise ratio in Kv3.3KO (blue) relative to WT (black). Data is presented as median and inter-quartiles. P values calculated using Mann-Whitney Rank Sum Test and statistically significant values displayed on each graph.

Temporal processing was tested by presenting suprathreshold sound stimuli at the neurons’ characteristic frequency in 6-8 month old mice and comparing WT and Kv3.3KO strains. MNTB neurons in WT responded with a phasic-tonic firing pattern (Fig. 7E,F) with short latencies (3.44ms [3.01; 4.21]; n=25; Fig. 7I,J) and minimal jitter (0.18ms [0.09; 0.51]; n=18; Fig. 7I,K). In contrast, Kv3.3KO neurons were slower to respond to sound with first spike latencies of 4.06ms [3.01; 4.21] (n=25; Mann-Whitney Rank Sum Test: *P*=0.015; Fig. 7I,J) and showed larger temporal variability (0.92ms [0.56; 1.50]; n=18; Mann-Whitney Rank Sum Test: *P*=0.001; Fig. 7I,K).

Kv3.3KO neurons lacked the onset phasic component in the peri-stimulus time histogram (Fig. 7F arrows). This inability to fire high instantaneous rates at sound onset was accompanied by a shift in the average inter-spike intervals from 2.16ms [1.48; 3.57] (n=22) in the WT to 3.10 [2.49; 4.19] (n=18) in the Kv3.3KO (Mann-Whitney Rank Sum Test: *P*≤0.001; Fig. 7G). Indeed, comparing peak firing rates during the first 10ms of the sound-evoked response revealed significantly lower rates in Kv3.3KOs (240Hz [180; 280]; n=18) compared to WTs (350Hz [200; 425]; n=22; Mann-Whitney Rank Sum Test: *P*=0.003; Fig. H). In contrast to the reduced sound-evoked firing rates, an increase in spontaneous firing was observed in the Kv3.3KOs (Kv3.3KO: 40Hz [19.3; 62.0]; n=19] compared to WT controls (16.5Hz [4.5; 42.8]; n=24; Mann-Whitney Rank Sum Test: *P*=0.044; Fig. 7L). Together, these combined changes caused a significant reduction in signal-to-noise ratio in the Kv3.3KO (2.77 [2.04; 3.99]; n=14) compared to WT controls (9.70 [4.38; 40.95]; n=18; Mann-Whitney Rank Sum Test: *P*=0.001; Fig. 7M).

## Discussion

Neurotransmitter release is triggered by calcium influx through voltage-gated calcium channels and is highly influenced by the presynaptic AP waveform. Deletion of Kv3.3 subunits increased presynaptic AP duration and transmitter release, enhancing short-term depression on repetitive stimulation even at relatively low frequencies. In contrast, deletion of Kv3.1 had little effect on the presynaptic AP or transmitter release. These observations were in agreement with a computational model of transmitter release in which Kv3.3 deletion increased vesicle release probability by 2-fold and to a lesser degree accelerated fast vesicle replenishment. This is consistent with increased calcium influx and activity-dependent facilitation of recycling. This enhanced EPSC depression in Kv3.3KOs impeded phasic AP firing during high frequency synaptic stimulation. *In vivo* recording showed that temporal fidelity and firing rates in the binaural auditory circuit were reduced in response to sound, whilst excitability and spontaneous AP firing increased in the Kv3.3KO. We conclude that presynaptic Kv3 channels require one (or more) Kv3.3 subunits to achieve fast and temporally precise transmitter release at this excitatory synapse.

Recent reports suggest that deletion of Kv3.3 inhibits both fast and slow endocytosis via non-ionic mechanisms, by altered interactions with the cytoskeleton (Wu ***et al.***, 2021). In contrast, we have measured accelerated recovery from short-term depression, along with increased presynaptic AP duration and transmitter release on deletion of Kv3.3 in mice of a similar age and at physiological temperatures. Our results are consistent with enhanced activity-dependent vesicle recycling (Wang & Kaczmarek, 1998) in the absence of Kv3.3, as a consequence of the increased AP duration and calcium influx (Yang ***et al.***, 2014). Our modelling indicates the mechanism is primarily an increase in vesicle release probability, but an acceleration of vesicle recycling also contributes to increased neurotransmitter release.

### Relevance to mechanisms of sound localization

Short duration APs permit high firing rates by minimising the absolute refractory period, thus Kv3 enhances well-timed high frequency AP trains and when input firing rates exceed the refractory limit, neurons expressing Kv3 ready switch to firing on alternate input spikes (Song ***et al.***, 2005). In this study we evoked MNTB neuron AP firing in response to presynaptic stimulation and asked how effective was the input/output across a range of firing frequencies from 100 to 600 Hz in each genotype. Although there was little difference at low firing frequencies, at the highest frequencies only the first few presynaptic APs generate postsynaptic APs with 1:1 fidelity (Phase I) before entering a chaotic phase (Phase II) with many failures, which resolves later in the train into precise firing but to alternate stimuli (Phase III). In the absence of Kv3.3 the MNTB firing passed into Phase II but did not converge into Phase III firing, consistent with the idea that presynaptic Kv3.3 enhances the stability of synaptic transmission.

Integration of information from both ears for the purpose of sound localization requires highly accurate transmission across the brainstem. This is facilitated by Kv3 channels within specific neurons, from cell bodies, axons and synaptic terminals, and across the network. *In vivo* extracellular recording from the MNTB measures simultaneously the presynaptic calyx and postsynaptic MNTB single unit APs. Accurate transmission of AP timing was compromised in the Kv3.3KO, and the peak firing rate was reduced. Additionally, spontaneous AP firing was elevated (with respect to WT animals), likely reflecting hyper-excitability upstream in the auditory pathway, with Kv3.3 being expressed in spiral ganglion (Kim ***et al.***, 2020) and the globular bushy cells (Li ***et al.***, 2001; Cao ***et al.***, 2007) (which give rise to the calyx of Held). The increased temporal jitter and AP delay further undermine interaural level discrimination (ILD) by reducing the gain and signal to noise and likely account for the impairment of sound localization in human spinocerebellar ataxia type 13 (SCA13) (Middlebrooks ***et al.***, 2013).

### Subunit composition

Knowledge of Kv channel subunit composition, interactions and precise location within identified neurons is key to understanding their extensive physiological roles in controlling neuronal excitability (Trimmer, 2015). Kv channel building is regulated by the N-terminal tetramerization domain (T1) (Li ***et al.***, 1992) to favour assembly of ‘dimers of dimers’ (Tu & Deutsch, 1999) usually from subunits within the same Kv family. Although early studies suggested otherwise, Kv3 subunits do not co-assemble with the accessory subunit gene families (Kv5, Kv6, Kv8 and Kv9) (Bocksteins ***et al.***, 2014), so Kv3 channels are likely composed of four Kv3 alpha subunits. Frequent co-expression of Kv3 subunits in the same neurons suggest functional channels exist as heteromers, indeed coimmunoprecipitation has revealed interactions of Kv3.1b with Kv3.4a and Kv3.2 in globus pallidus neurons {Baranauskas:2003fw, (Hernandez-Pineda ***et al.***, 1999), and with Kv3.3 in the cerebellum (Chang ***et al.***, 2007a). The data here however, suggests that presynaptic Kv3 channels at the calyx of Held require an obligatory Kv3.3 subunit, since TEA (which will block all Kv3 heteromers) showed little effect on transmission in the Kv3.3KO. This result also implies that Kv3.1 (and Kv3.2 or Kv3.4) do not have a prominent independent role in the terminal and while heteromers with Kv3.3 may occur, Kv3.1 is certainly not necessary for Kv3.3 localisation to the presynaptic membrane.

### Do interactions with the cytoskeleton dictate subcellular localisation?

Proximity is a powerful influence on the voltage control of other ion channels by K^+^ channels, and hence the permutations and complexity of channel subunit assembly enables optimal localization (Trimmer, 2015) through trafficking, insertion and stabilization. It has been argued that Kv3.3 has a non-ionic role in modulating endocytosis, but given the evidence for linkage to the actin cytoskeleton (Zhang ***et al.***, 2016) and of Kv3.1b to the extracellular matrix (Stevens ***et al.***, 2021), a scaffolding interaction is also a reasonable hypothesis. Insertion of potassium channels into key sites of excitability (axon initial segments, nodes of Ranvier, heminodes and synaptic terminals) permits tight control of voltage to avoid hyper-excitability. An ionic hypothesis for the role of Kv3 is that insertion into presynaptic locations facilitates control of voltage-gated sodium and calcium channels, and thereby contributes to enabling recovery from inactivation and refining Ca^2+^ influx in adaptation of transmitter release from a microdomain to a nanodomain architecture (Meinrenken ***et al.***, 2003) in neurons possessing presynaptic Kv3.3.

### Kv3.3 dysfunction and disease

Immunohistochemical studies have localized Kv3.3 subunits to many different synaptic terminals across the brain, from spiral ganglion afferent processes (Kim ***et al.***, 2020), medial vestibular nuclei (Brooke ***et al.***, 2010), cerebellar dentate nucleus (Alonso-Espinaco ***et al.***, 2008), parallel fibre synapses (Puente ***et al.***, 2010), posterior thalamic nucleus (Chang ***et al.***, 2007b) and the neuromuscular junction (Brooke ***et al.***, 2004). The current study suggests that mutations associated with Kv3.3 such as SCA13, cause a range of neurological defects some of which will reflect aberrant neurotransmitter release. It is certainly the case that in SCA13 mutation R420H (Middlebrooks ***et al.***, 2013), the observations made here support the hypothesis that SCA13 disrupts the physiological process of interaural level discrimination, and thereby undermines sound source localization (Tollin, 2003).

The combinatorial potential of Kv3 channel subunits gives rise to a spectrum of physiological roles in fast-spiking neurons (Choudhury ***et al.***, 2020) and interneurons (Chang ***et al.***, 2007a). This study shows that presynaptic Kv3.3 subunits further enhance the biophysical ‘speed limit’ for information transmission at those synapses where it is expressed.

## METHODS

Experiments were conducted in accordance with the Animals (Scientific Procedures) Act UK 1986 and as revised by the European Directive 2010/63/EU on the protection of animals used for scientific purposes. All procedures were approved by national oversight bodies (UK Home Office, or Bavarian district government, ROB-55.2-2532.Vet_02-18-1183) and the local animal research ethics review committees.

Experiments were conducted on CBA/Crl mice (wildtype, WT) and knockout were mice backcrossed for 10 generations onto this CBA/Crl background (Choudhury ***et al.***, 2020). PCR genotyping was done from ear notch samples made at P10. Mice were housed and breeding colonies maintained at the preclinical research facility (PRF) at the University of Leicester, subject to a normal 12hr light/dark cycle and with free access to food and water (*ad libitum*). Both male and female animals were used in experiments with ages ranging from P10-25 for electrophysiology to 6 months of age for *in vivo* and auditory brainstem response (ABR) recordings.

### mRNA sequencing

Mice were killed by decapitation and brainstems were removed into ‘RNA Later’ stabilization solution (Invitrogen, Cat# AM7020), before dissection to isolate both cochlear nuclei. Tissue from 3 mice CBA/Crl mice were pooled into 9 individual samples (27 mice total) before phenol extraction to isolate RNA. RNA purity, integrity, and concentration was assessed by UV-Vis spectroscopy (Nanodrop 8000) and capillary electrophoresis (Agilent Bioanalyzer 2000). Samples with RIN (RNA integrity) < 7 were discarded. cDNA libraries were constructed using the NEBNext® Ultra™ Directional RNA Library Prep Kit for Illumina® sequencing performed using the Illumina NextSeq500 High Output (v2, 150 cycles) kit and the Illumina® NextSeq500. Analysis of sequencing data was performed using Illumina Basespace. Using FastQC toolkit (Babraham Bioinformatics) total reads were trimmed of low quality reads (Q<20), poly-A/T tails >10bp, and adapter sequences before alignment to the mm10 (GRCM387) mouse genome (Ensembl) using TopHat2. Output values represented as Fragments per kilobase of transcript per million mapped reads (FPKM).

### Electrophysiology

#### *In vitro* brain slice preparation

Mice were killed by decapitation, the brainstem removed and placed into ice-cold artificial cerebrospinal fluid aCSF, oxygenated with 95%O_2_/5%CO_2_, containing (in mM): sucrose (250), KCl (2.5), NaHCO_3_ (26), NaH_2_PO_4_ (1.25), D-Glucose (10), ascorbic acid (0.5) MgCl_2_ (4), and CaCl_2_ (0.1). For presynaptic recordings 100 μm thick transverse slices or 250 μm thick slices for postsynaptic recordings were prepared in a pre-cooled chamber using a Leica VT1200S vibratome. Slices were allowed to recover for 1 hour at 37°C in normal aCSF (Choudhury ***et al.***, 2020), continually bubbled with 95%O_2_/5%CO_2_ and subsequently allowed to passively cool to room temperature. The aCSF (310 mOsm) contained (in mM): NaCl (125), NaHCO_3_ (26), D-Glucose (10), KCl (2.5), myo-inositol (3), NaH_2_PO_4_ (1.25), sodium pyruvate (1), ascorbic acid (0.5), MgCl_2_ (1), and CaCl_2_ (2).

### Presynaptic recordings

Mice aged P10-P12 were used for presynaptic calyx recordings. For each experiment, slices were placed in a recording chamber of a Nikon E600FN upright microscope and cells visualised with a 60x DIC water-immersion objective (Lucas ***et al.***, 2018). Slices were continuously perfused with normal aCSF saturated with 95%O_2_/5%CO_2_, (as above), heated to 35°C ± 1, at a rate of 1ml/min. Whole-cell patch recordings were made using thick-walled borosilicate capillaries (1.5 mm OD, 0.86 mm ID) with a resistance of 4-6MΩ, filled with an internal solution composed of (in mM): KGluconate (97.5), KCl (32.5), HEPES (40), EGTA (0.2), MgCl_2_ (1), K_2_ATP (2.2), Na_2_GTP (0.3), pH adjusted to 7.2 with KOH (295 mOsm).

Recordings were made with a Multiclamp 700A amplifier (Molecular Devices), 1322A digidata (Axon Instruments) and Pclamp 10 software (Molecular Devices) for acquisition and analysis. Electrode and cell capacitance were compensated and series resistances were corrected for with recordings discarded when series resistances reached >20MΩ before compensation. All recordings were compensated by 70%. Signals were digitised at 100kHz and filtered at 10kHz. The stated voltages were not corrected for a liquid junction potential of 11mV.

Current-voltage relationships were generated in voltage-clamp over a range of command voltages from −110mV −+30mV in 10mV incremental steps. Steps were 150ms in duration and separated by 1s intervals. Action potentials measured in current-clamp were generated using short depolarising current steps (50pA increments) of 50ms duration, the first (threshold) evoked action potential was analysed. Pipette capacitance was neutralised and the bridge balanced. Hyperpolarising current steps (−50pA, 150ms) were used to determine membrane resistance. For both voltage and current clamp experiments, voltage or current commands were from −70mV.

### Postsynaptic recordings

The same experimental setup as described above was used for postsynaptic recordings. Borosilicate glass pipettes (2.5-3.5 MΩ resistance) were filled with a solution containing (in mM) KGluconate (120), KCl (10), HEPES (40), EGTA (0.2), MgCl2 (1), K2ATP (2.2). Electrode and cell capacitance were compensated and series resistances were corrected, with recordings discarded when series resistances reached >10MΩ (before compensation) or changed by >10% during the recording. All recordings were compensated by 70%, signals were digitised at 100kHz and filtered at 10kHz.

Mice aged P20-P27 were used for studying synaptic physiology. Axons giving rise to the calyx terminal were stimulated using a concentric bipolar electrode (FHC, inc #CBAD75S) placed at the midline of a brainstem slice, controlled by a constant voltage stimulator box (DS2A, Digitimer) triggered by the Pclamp 10 software. Axons were subjected to low-frequency stimulation (0.3Hz) in order to determine the threshold for generating evoked excitatory postsynaptic currents (EPSCs) after which stimulation trains of 5 x 100, (separated by 20s intervals), 200 and 600Hz for 800ms separated by 30s intervals were applied. Time was allowed for the internal patch solution to equilibrate (5 minutes) before stimulation trains were applied. EPSC recordings were conducted from a holding potential of −40mV (to inactivate voltage-gated sodium channels) and inhibitory transmission blocked by adding 0.5μM strychnine hydrochloride to external aCSF.

### Postsynaptic recordings of action potential trains at high frequencies

Current clamp recordings were made from MNTB neurons stimulated presynaptically with a bipolar concentric electrode (FHC, inc #CBAD75S). Trains of 0.1ms stimuli of variable voltages (3-15V) were delivered to the presynaptic axon at 100, 200, 300, 400 and 600 Hz using a constant voltage stimulation box (DS2A, Digitimer) triggered by the pClamp10 software. Evoked responses were recorded from the postsynaptic neuron under current clamp, with resting membrane potentials adjusted to −60 mV. Stimulation trains of the duration of 800ms at the different frequencies were repeated 3 times and separated by 20s intervals to allow for recovery.

### Voltage and current-clamp analysis

Electrophysiology analysis was conducted using Clampfit 10 software (Molecular Devices). Current amplitudes were measured as the steady-state current towards the end of the 150ms voltage step. Presynaptic action potentials were analyzed using the threshold detection function. The threshold was set to the voltage of action potential activation and the relative amplitude defined as the difference between voltage at the peak and voltage at the threshold. Half-width is defined as the time delay between upstroke and downstroke at half-maximal amplitude. Rise and decay slopes were measured from 10-90% of peak amplitude.

Single excitatory postsynaptic currents were analyzed using the threshold detection function in Pclamp. Baseline was defined as the resting current before stimulation. The threshold for detection was set to twice the standard deviation of the noise level. Peak amplitudes were defined as the difference between the current and the peak and the current at the baseline. Rise times were measured from 10-90% of the peak and decay times were measured from 90% of the peak to 1s after the peak. Charge was measured as the area under the curve, between the peak and baseline.

EPSC trains were analyzed by normalizing each response to the first response in the train then fitting a single exponential to normalized amplitudes of responses and extracting the decay tau and steady-state amplitudes of the exponential fit to define the rate at which responses underwent short-term depression and the extent to which they depressed.

#### *In vivo* physiology

Adult (6-8 month) Kv3.3 knockout mice of either sex (n=5) and five age-matched CBA wild type mice were anesthetized with a subcutaneous injection of 0.01ml/g MMF (0.5mg/kg body weight Medetomidine, 5.0mg/kg body weight Midazolam and 0.05mg/kg body weight Fentanyl). They were placed on a temperature-controlled heating pad (WPI: ATC 1000) in a soundproof chamber (Industrial Acoustics). Depth of anaesthesia was measured using the toe pinch reflex and animals responding were given supplemental MMF at 1/3 the initial dose. The mice were stabilized in a custom stereotaxic device. An incision was made at the top of the skull, followed by a craniotomy just anterior to the lambda suture intersection. The skull was tilted to provide access to the auditory brainstem. A ground electrode was placed in the muscle at the base of the neck. Glass microelectrodes were pulled from glass capillaries so that the resistance was 5-20 MW when filled with 3M KCl solution. Signals were amplified (AM Systems, Neuroprobe 1600), filtered (300-3000Hz; Tucker-Davis-Technologies PC1) and recorded (~50 kHz sampling rate) with a Fireface UFX audio interface (RME). AudioSpike software (HoerrTech) was used to calibrate the multi-field magnetic speakers, generate stimuli and record action potentials. Stimuli consisted of pure tones (50-100ms duration, 5ms rise/fall time) at varying intensity (0-90dB SPL) and were presented through hollow ear bars connected to the speakers with Tygon tubing. PSTHs were assessed at characteristic frequency (CF) and 80dB SPL. MNTB neurons were identified by their excitatory response to contralateral sound stimulation and their typical complex waveform (Kopp-Scheinpflug ***et al.***, 2003), consisting of a presynaptic potential (preAP), a synaptic delay (SD) and a postsynaptic potential (AP).

### Auditory-evoked brainstem response

ABR equipment set-up and recordings have previously been described in detail in (Ingham ***et al.***, 2011). Briefly, mice were anesthetized with fentanyl (0.04 mg/kg), midazolam (4 mg/kg), and medetomidine (0.4 mg/kg) by intraperitoneal injection. Animals were placed on a heated mat inside a sound-attenuated chamber, and electrodes were inserted sub-dermally; below the right pinnae, into the muscle mass below the left ear, and at the cranial vertex. ABR responses were collected, amplified, and averaged using the TDT System3 (Tucker Davies Technology) in conjunction with custom ‘Averager’ software, provided by the Wellcome Trust Sanger Institute. Binaural stimuli were delivered in the form of a 0.1 ms broadband click. All stimuli were presented in 5 dB SPL rising steps to 95 dB SPL, and responses were averaged 512 times per step. Recordings were averaged over a 20ms period with a 300-3000Hz bandwidth filter and a gain of 25000x. Wave amplitude and latencies were analyzed using the Auditory Wave Analysis Python script developed by Bradley Buran (Eaton-Peabody Laboratory), and calculated as the difference between peak and valley (μV) and time to wave peak (s), respectively.

### Computation Model

A simple model with activity-dependent vesicle recycling (Graham ***et al.***, 2004; Billups ***et al.***, 2005; Lucas ***et al.***, 2018) was used. In the model, vesicles in a releasable pool of normalised size *n* may release with a fixed probability *p*=*P*_*v*_ on the arrival of a presynaptic action potential at time *s* to give an EPSC amplitude proportional to *np* (equ. 3). Vesicles in this releasable pool are replenished up to the maximum normalised pool size of n=1 at a rate τ_*r*_ from an infinite reserve pool (equ. 1). In the absence of presynaptic action potentials, replenishment proceeds at a constant background rate (time constant τ_*b*_). Following a presynaptic action potential, the replenishment rate is instantaneously raised to a higher rate, τ_*h*_ (equ. 3) which then decays back to the background rate with time constant τ_*d*_ (equ. 2a). The model equations are:

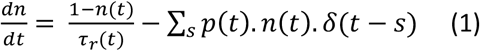

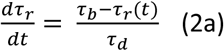

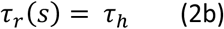

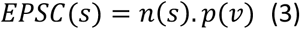

The model is implemented in Matlab. Differential equations are solved by simple forward Euler integration.

### Statistics

Statistical analysis of the *in vitro* data was performed in Graphpad Prism V7 unless otherwise specified. Data were tested for a normal gaussian distribution using a Shapiro-Wilk normality test and parametric (one-way ANOVA) or non-parametric tests (Kruksal-Wallis ANOVA) applied as appropriate. Multiple comparisons were corrected for using Tukey‘s multiple comparisons test or Dunn’s multiple comparison test post-hoc and a P value of <0.05 was taken as significant. The statistical tests applied are noted in the figure legends and corresponding text. Data is represented as mean ± SD unless otherwise stated. *In vivo* data are presented as medians and inter-quartiles in text numbers and figures in addition to individual data points. Statistical analyses of the *in vivo* data were performed with SigmaStat/SigmaPlot™. Normality was tested by the Shapiro-Wilk Test. Comparisons between data sets were made using parametric tests for normally distributed data (two-tailed Student’s t-test for comparing two groups) and when the normality assumption was violated, a non-parametric test (Mann-Whitney Rank Sum Test) was used.

## Acknowledgements

We are grateful to the Preclinical Research Facility at the University of Leicester for the animal care, husbandry and expert assistance provided, also to Neil Ingram for assistance and advice in setting up the ABR recording system. This research was funded by a BBSRC project grant (IDF) and a BBSRC Case PhD Studentship (AR, NP, IDF) including support from Autifony Therapeutics Ltd; further funding was provided by DFG SFB870 A-10 (CKS). We are grateful for the support provided to IDF by Benedikt Grothe during a sabbatical in the Division of Neurobiology, Faculty of Biology Ludwig Maximilian University, Munich, Germany.

---

**Figure S1:**
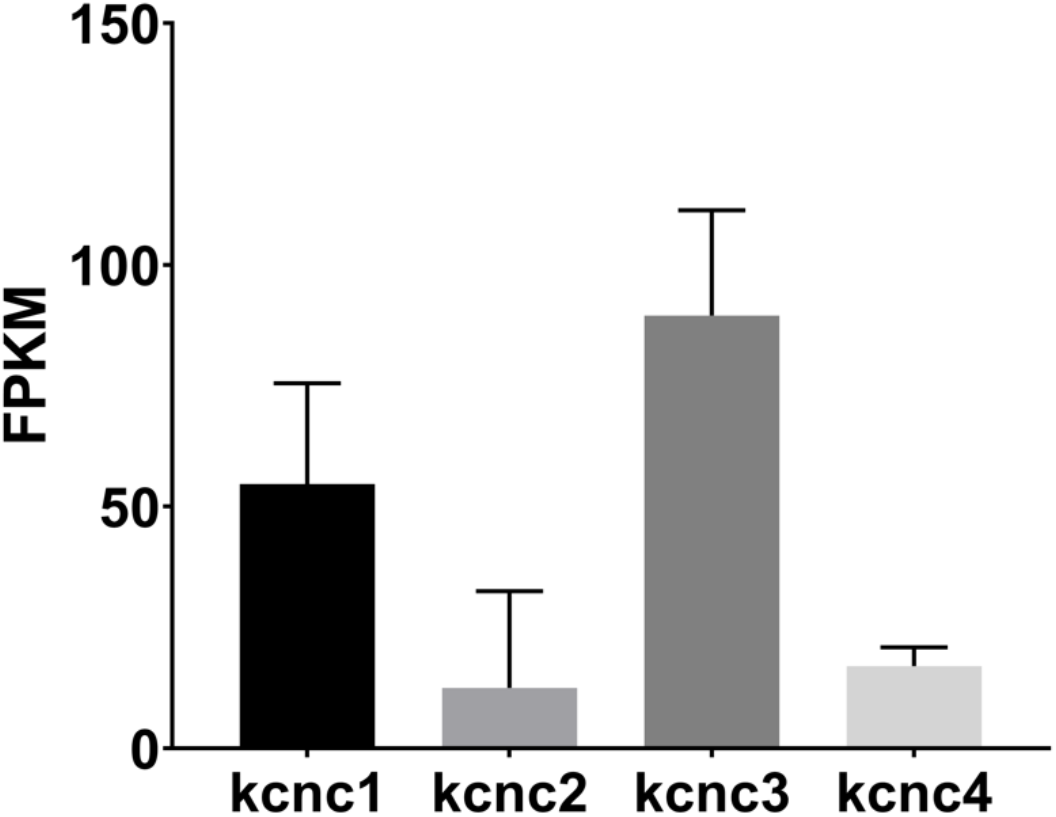
WT mRNA Levels for kcnc family genes in the cochlear nucleus which show a similar pattern of expression to those measured previously for the MNTB (Choudhury et al., 2020). The data shows predominance of kcnc1 (Kv3.1) and kcnc3 (Kv3.3). Mean ± SD is plotted for 9 measured samples, with each sample containing 3 cochlear nuclei. Data is presented as **Fragments per kilobase of transcript per million mapped reads (FPKM)**.

## REFERENCES

Alonso-Espinaco V, Elezgarai I, Díez-García J, Puente N, Knöpfel T & Grandes P (2008). Subcellular localization of the voltage-gated potassium channels Kv3.1b and Kv3.3 in the cerebellar dentate nucleus of glutamic acid decarboxylase 67-green fluorescent protein transgenic mice. Neuroscience 155, 1059–1069.

Beiderbeck B, Myoga MH, Müller NIC, Callan AR, Friauf E, Grothe B & Pecka M (2018). Precisely timed inhibition facilitates action potential firing for spatial coding in the auditory brainstem. Nature Communications 9, 1771.

Billups B, Graham BP, Wong A & Forsythe ID (2005). Unmasking group III metabotropic glutamate autoreceptor function at excitatory synapses in the rat CNS. J Physiol (Lond) 565, 885–896.

Bocksteins E, Mayeur E, Van Tilborg A, Regnier G, Timmermans J-P & Snyders DJ (2014). The Subfamily-Specific Interaction between Kv2.1 and Kv6.4 Subunits Is Determined by Interactions between the N- and C-termini ed. Phillips W. PLoS ONE; DOI: 10.1371/journal.pone.0098960.

Borst JG & Sakmann B (1998). Calcium current during a single action potential in a large presynaptic terminal of the rat brainstem. J Physiol (Lond) 506 (Pt 1), 143–157.

Brew HM & Forsythe ID (1995). Two voltage-dependent K+ conductances with complementary functions in postsynaptic integration at a central auditory synapse. J Neurosci 15, 8011–8022.

Brooke RE, Corns L, Edwards IJ & Deuchars J (2010). Kv3.3 immunoreactivity in the vestibular nuclear complex of the rat with focus on the medial vestibular nucleus: targeting of Kv3.3 neurones by terminals positive for vesicular glutamate transporter 1. Brain Res 1345, 45–58.

Brooke RE, Moores TS, Morris NP, Parson SH & Deuchars J (2004). Kv3 voltage-gated potassium channels regulate neurotransmitter release from mouse motor nerve terminals. Eur J Neurosci 20, 3313–3321.

Cao X-J, Shatadal S & Oertel D (2007). Voltage-sensitive conductances of bushy cells of the Mammalian ventral cochlear nucleus. J Neurophysiol 97, 3961–3975.

Chang SY, Zagha E, Kwon ES, Ozaita A, Bobik M, Martone ME, Ellisman MH, Heintz N & Rudy B (2007a). Distribution of Kv3.3 potassium channel subunits in distinct neuronal populations of mouse brain. J Comp Neurol 502, 953–972.

Chang SY, Zagha E, Kwon ES, Ozaita A, Bobik M, Martone ME, Ellisman MH, Heintz N & Rudy B (2007b). Distribution of Kv3.3 potassium channel subunits in distinct neuronal populations of mouse brain. J Comp Neurol 502, 953–972.

Choudhury N, Linley D, Richardson A, Anderson M, Robinson SW, Marra V, Ciampani V, Walter SM, Kopp-Scheinpflug C, Steinert JR & Forsythe ID (2020). Kv3.1 and Kv3.3 subunits differentially contribute to Kv3 channels and action potential repolarization in principal neurons of the auditory brainstem. J Physiol (Lond) 598, 2199–2222.

Coetzee WA, Amarillo Y, Chiu J, Chow A, Lau D, McCormack T, Moreno H, Nadal MS, Ozaita A, Pountney D, Saganich M, Vega-Saenz de Miera E & Rudy B (1999). Molecular diversity of K+ channels. Ann N Y Acad Sci 868, 233–285.

Devaux J, Alcaraz G, Grinspan J, Bennett V, Joho R, Crest M & Scherer SS (2003). Kv3.1b is a novel component of CNS nodes. Journal of Neuroscience 23, 4509–4518.

Du J, Haak LL, Phillips-Tansey E, Russell JT & McBain CJ (2000). Frequency-dependent regulation of rat hippocampal somato-dendritic excitability by the K+ channel subunit Kv2.1. J Physiol (Lond) 522 Pt 1, 19–31.

Espinosa F, McMahon A, Chan E, Wang S, Ho CS, Heintz N & Joho RH (2001). Alcohol hypersensitivity, increased locomotion, and spontaneous myoclonus in mice lacking the potassium channels Kv3.1 and Kv3.3. J Neurosci 21, 6657–6665.

Espinosa F, Torres-Vega MA, Marks GA & Joho RH (2008). Ablation of Kv3.1 and Kv3.3 potassium channels disrupts thalamocortical oscillations in vitro and in vivo. Journal of Neuroscience 28, 5570–5581.

Forsythe ID (1994). Direct patch recording from identified presynaptic terminals mediating glutamatergic EPSCs in the rat CNS, in vitro. J Physiol (Lond) 479 (Pt 3), 381–387.

Forsythe ID, Tsujimoto T, Barnes-Davies M, Cuttle MF & Takahashi T (1998). Inactivation of Presynaptic Calcium Current Contributes to Synaptic Depression at a Fast Central Synapse. Neuron 20, 11–11.

Geiger JR & Jonas P (2000). Dynamic control of presynaptic Ca(2+) inflow by fast-inactivating K(+) channels in hippocampal mossy fiber boutons. Neuron 28, 927–939.

Graham BP, Wong A & Forsythe ID (2004). A multi-component model of depression at the calyx of Held. Neurocomputing 58, 449–454.

Grissmer S, Nguyen AN, Aiyar J, Hanson DC, Mather RJ, Gutman GA, Karmilowicz MJ, Auperin DD & Chandy KG (1994). Pharmacological characterization of five cloned voltage-gated K+ channels, types Kv1.1, 1.2, 1.3, 1.5, and 3.1, stably expressed in mammalian cell lines. Molecular Pharmacology 45, 1227–1234.

Hennig MH, Postlethwaite M, Forsythe ID & Graham BP (2008). Interactions between multiple sources of short-term plasticity during evoked and spontaneous activity at the rat calyx of Held. J Physiol (Lond) 586, 3129–3146.

Hernandez-Pineda R, Chow A, Amarillo Y, Moreno H, Saganich M, Vega-Saenz de Miera EC, Hernandez-Cruz A & Rudy B (1999). Kv3.1-Kv3.2 channels underlie a high-voltage-activating component of the delayed rectifier K+ current in projecting neurons from the globus pallidus. J Neurophysiol 82, 1512–1528.

Ingham NJ, Pearson S & Steel KP (2011). Using the Auditory Brainstem Response (ABR) to Determine Sensitivity of Hearing in Mutant Mice. Curr Protoc Mouse Biol 1, 279–287.

Johnston J, Forsythe ID & Kopp-Scheinpflug C (2010). Going native: voltage-gated potassium channels controlling neuronal excitability. J Physiol (Lond) 588, 3187–3200.

Joho RH, Ho CS & Marks GA (1999). Increased gamma- and decreased delta-oscillations in a mouse deficient for a potassium channel expressed in fast-spiking interneurons. J Neurophysiol 82, 1855–1864.

Joho RH, Street C, Matsushita S & Knöpfel T (2006). Behavioral motor dysfunction in Kv3-type potassium channel-deficient mice. Genes Brain Behav 5, 472–482.

Joris PX & Trussell LO (2018). The Calyx of Held: A Hypothesis on the Need for Reliable Timing in an Intensity-Difference Encoder. Neuron 100, 534–549.

Kaczmarek LK & Zhang Y (2017). Kv3 Channels: Enablers of Rapid Firing, Neurotransmitter Release, and Neuronal Endurance. Physiological Reviews 97, 1431–1468.

Kim WB, Kang K-W, Sharma K & Yi E (2020). Distribution of Kv3 Subunits in Cochlear Afferent and Efferent Nerve Fibers Implies Distinct Role in Auditory Processing. Exp Neurobiol 29, 344–355.

Kopp-Scheinpflug C, Lippe WR, Dörrscheidt GJ & Rübsamen R (2003). The medial nucleus of the trapezoid body in the gerbil is more than a relay: comparison of pre- and postsynaptic activity. J Assoc Res Otolaryngol 4, 1–23.

Kopp-Scheinpflug C, Tolnai S, Malmierca MS & Rübsamen R (2008). The medial nucleus of the trapezoid body: Comparative physiology. Neuroscience 154, 160–170.

Labro AJ, Priest MF, Lacroix JJ, Snyders DJ & Bezanilla F (2015). Kv3.1 uses a timely resurgent K(+) current to secure action potential repolarization. Nature Communications 6, 10173.

Li M, Jan YN & Jan LY (1992). Specification of subunit assembly by the hydrophilic amino-terminal domain of the Shaker potassium channel. Science 257, 1225–1230.

Li W, Kaczmarek LK & Perney TM (2001). Localization of two high-threshold potassium channel subunits in the rat central auditory system. J Comp Neurol 437, 196–218.

Lien CC & Jonas P (2003). Kv3 potassium conductance is necessary and kinetically optimized for high-frequency action potential generation in hippocampal interneurons. J Neurosci 23, 2058–2068.

Lucas SJ, Michel CB, Marra V, Smalley JL, Hennig MH, Graham BP & Forsythe ID (2018). Glucose and lactate as metabolic constraints on presynaptic transmission at an excitatory synapse. J Physiol (Lond) 596, 1699–1721.

Meinrenken CJ, Borst JGG & Sakmann B (2003). Local routes revisited: the space and time dependence of the Ca2+ signal for phasic transmitter release at the rat calyx of Held. J Physiol (Lond) 547, 665–689.

Middlebrooks JC, Nick HS, Subramony SH, Advincula J, Rosales RL, Lee LV, Ashizawa T & Waters MF (2013). Mutation in the kv3.3 voltage-gated potassium channel causing spinocerebellar ataxia 13 disrupts sound-localization mechanisms. PLoS ONE 8, e76749.

Neher E (2017). Some Subtle Lessons from the Calyx of Held Synapse. Biophys J 112, 215–223.

Puente N, Mendizabal-Zubiaga J, Elezgarai I, Reguero L, Buceta I & Grandes P (2010). Precise localization of the voltage-gated potassium channel subunits Kv3.1b and Kv3.3 revealed in the molecular layer of the rat cerebellar cortex by a pre-embedding immunogold method. Histochem Cell Biol 134, 403–409.

Ritzau-Jost A, Tsintsadze T, Krueger M, Ader J, Bechmann I, Eilers J, Barbour B, Smith SM & Hallermann S (2021). Large, Stable Spikes Exhibit Differential Broadening in Excitatory and Inhibitory Neocortical Boutons. Cell Rep 34, 108612.

Rowan MJM, Tranquil E & Christie JM (2014). Distinct kv channel subtypes contribute to differences in spike signaling properties in the axon initial segment and presynaptic boutons of cerebellar interneurons. Journal of Neuroscience 34, 6611–6623.

Rudy B & McBain CJ (2001). Kv3 channels: voltage-gated K+ channels designed for high-frequency repetitive firing. Trends in Neurosciences 24, 517–526.

Rudy B, Chow A, Lau D, Amarillo Y, Ozaita A, Saganich M, Moreno H, Nadal MS, Pinedar RH, Cruz AH, Erisir A, Leonard C & Vega-Saenz de Miera E (2002). Contributions of Kv3 channels to neuronal excitability. Ann N Y Acad Sci 868, 304–343.

Sakaba T & Neher E (2003). Involvement of actin polymerization in vesicle recruitment at the calyx of Held synapse. J Neurosci 23, 837–846.

Schneggenburger R & Forsythe ID (2006). The calyx of Held. Cell Tissue Res 326, 311–337.

Song P, Yang Y, Barnes-Davies M, Bhattacharjee A, Hamann M, Forsythe ID, Oliver DL & Kaczmarek LK (2005). Acoustic environment determines phosphorylation state of the Kv3.1 potassium channel in auditory neurons. Nat Neurosci 8, 1335–1342.

Steinert JR, Robinson SW, Tong H, Haustein MD, Kopp-Scheinpflug C & Forsythe ID (2011). Nitric oxide is an activity-dependent regulator of target neuron intrinsic excitability. Neuron 71, 291–305.

Stevens SR, Longley CM, Ogawa Y, Teliska LH, Arumanayagam AS, Nair S, Oses-Prieto JA, Burlingame AL, Cykowski MD, Xue M & Rasband MN (2021). Ankyrin-R regulates fast-spiking interneuron excitability through perineuronal nets and Kv3.1b K+ channels. Elife; DOI: 10.7554/eLife.66491.

Taschenberger H & Gersdorff von H (2000). Fine-tuning an auditory synapse for speed and fidelity: developmental changes in presynaptic waveform, EPSC kinetics, and synaptic plasticity. J Neurosci 20, 9162–9173.

Taschenberger H, Leao RM, Rowland KC, Spirou GA & Gersdorff von H (2002). Optimizing synaptic architecture and efficiency for high-frequency transmission. Neuron 36, 1127–1143.

Tollin DJ (2003). The lateral superior olive: a functional role in sound source localization. The Neuroscientist 9, 127–143.

Trimmer JS (2015). Subcellular localization of K+ channels in mammalian brain neurons: remarkable precision in the midst of extraordinary complexity. Neuron 85, 238–256.

Tu LW & Deutsch C (1999). Evidence for dimerization of dimers in K+ channel assembly. Biophys J 76, 2004–2017.

Wang LY & Kaczmarek LK (1998). High-frequency firing helps replenish the readily releasable pool of synaptic vesicles. Nature 394, 384–388.

Wang LY, Gan L, Forsythe ID & Kaczmarek LK (1998). Contribution of the Kv3.1 potassium channel to high-frequency firing in mouse auditory neurones. J Physiol (Lond) 509, 183–194.

Weiser M, Vega-Saenz de Miera E, Kentros C, Moreno H, Franzen L, Hillman D, Baker H & Rudy B (1994). Differential expression of Shaw-related K+ channels in the rat central nervous system. J Neurosci 14, 949–972.

Wong A, Graham BP, Billups B & Forsythe ID (2003). Distinguishing between presynaptic and postsynaptic mechanisms of short-term depression during action potential trains. J Neurosci 23, 4868–4877.

Wu X-S, Subramanian S, Zhang Y, Shi B, Xia J, Li T, Guo X, El-Hassar L, Szigeti-Buck K, Henao-Mejia J, Flavell RA, Horvath TL, Jonas EA, Kaczmarek LK & Wu L-G (2021). Presynaptic Kv3 channels are required for fast and slow endocytosis of synaptic vesicles. Neuron 92, 8328.

Yang Y-M, Wang W, Fedchyshyn MJ, Zhou Z, Ding J & Wang L-Y (2014). Enhancing the fidelity of neurotransmission by activity-dependent facilitation of presynaptic potassium currents. Nature Communications 5, 4564.

Young SM & Veeraraghavan P (2021). Presynaptic voltage-gated calcium channels in the auditory brainstem. Mol Cell Neurosci 112, 103609.

Zagha E, Manita S, Ross WN & Rudy B (2010). Dendritic Kv3.3 Potassium Channels in Cerebellar Purkinje Cells Regulate Generation and Spatial Dynamics of Dendritic Ca2+ Spikes. J Neurophysiol 103, 3516–3525.

Zhang Y, Zhang X-F, Fleming MR, Amiri A, El-Hassar L, Surguchev AA, Hyland C, Jenkins DP, Desai R, Brown MR, Gazula V-R, Waters MF, Large CH, Horvath TL, Navaratnam D, Vaccarino FM, Forscher P & Kaczmarek LK (2016). Kv3.3 Channels Bind Hax-1 and Arp2/3 to Assemble a Stable Local Actin Network that Regulates Channel Gating. Cell 165, 434–448.

